# Compositional pretraining improves computational efficiency and matches animal behavior on complex tasks

**DOI:** 10.1101/2024.01.12.575461

**Authors:** David Hocker, Christine M. Constantinople, Cristina Savin

**Affiliations:** Center for Neural Science, New York University, New York, NY 10003, USA; Center for Data Science, New York University, New York, NY 10011, USA

**Author notes:** Co-senior authors (Christine M. Constantinople, Cristina Savin).

## Abstract

Recurrent neural networks (RNN) are ubiquitously used in neuroscience to capture both neural dynamics and behaviors of living systems. However, when it comes to complex cognitive tasks, training RNNs with traditional methods can prove difficult and fall short of capturing crucial aspects of animal behavior. Here we propose a principled approach for identifying and incorporating compositional tasks as part of RNN training. Taking as target a temporal wagering task previously studied in rats, we design a pretraining curriculum of simpler cognitive tasks that reflect relevant sub-computations. We show that this pretraining substantially improves learning efficacy and is critical for RNNs to adopt similar strategies as rats, including long-timescale inference of latent states, which conventional pretraining approaches fail to capture. Mechanistically, our pretraining supports the development of slow dynamical systems features needed for implementing both inference and value-based decision making. Overall, our approach is an important step for endowing RNNs with relevant inductive biases, which is important when modeling complex behaviors that rely on multiple cognitive computations.

## 2 Introduction

Recurrent neural networks (RNN) are frequently used in neuroscience to study the role of coordinated network activity in supporting trained behaviors, with the hope of generating hypotheses about neural mechanisms that can be tested in animal models [1]. As cognitive tasks increase in complexity, training RNNs to mimic animals becomes increasingly difficult. First, RNNs may fail to learn the task, especially in reinforcement learning settings with salient suboptimal solutions, sparse rewards, or long temporal dependencies [2]. Second, the solutions found by RNNs trained on complex tasks can fail to reflect the computational strategies employed by animals, limiting their utility as models for neuroscience.

For difficult tasks, machine learning has conventionally used *pretraining*, a process in which neural networks are first trained on variants of the target task, to help improve final target task performance. This could involve slowly increasing the complexity along a particular dimension (e.g., gradually increasing the delay in a task requiring working memory), as a form of curriculum learning [3, 4]. Alternatively, pretraining could involve families of related tasks that require harnessing shared structure to bolster learning, as in meta-learning [5–7] (e.g., rule learning as a biological analogy). Importantly, such approaches usually train within the context of the target task, without taking into account broader computational goals or inductive biases.

Training animals to perform cognitive tasks presents its own challenges: complex behaviors require long training times and animals may adopt suboptimal strategies. To address this, behavioral scientists have developed their own pretraining procedures, in the form of *behavioral shaping* [8]. Shaping is the process by which an animal is gradually exposed to (and thus learns) the core components of a behavioral task, such as whether or not to interact with effectors like levers or ports, or associate sensory cues with rewards or abstract rules, based on reinforcement. Shaping is ubiquitous in behavioral neuroscience and often decomposes the target task into functional sub-elements (e.g. pressing a lever can provide reward); as with artificial agent pretraining, complexity is gradually increased by incorporating additional elements (e.g. sensory defined context). Over the training stages, the animal learns how to combine each sub-task in an appropriate way (e.g. lever needs to be pressed only when light is on). Nonetheless, despite its practical utility in increasing learning efficiency in animals, modeling these training-time learning experiences during RNN training remains uncommon [9].

A third, largely ignored, form of meaningful pretraining is the experience that the animals bring with them to the experiment in the form of *inductive biases*. Subjects learn many cognitive tasks throughout their lifetime, and previously-learned tasks can impact their behavior in new ones [10–12]. Many factors including training history [13], early experiences [14–17] or inductive biases established by evolution [18] can influence learning outcomes and corresponding behavioral strategies. Therefore, modeling relevant prior experiences may increase the ability of RNNs to capture complex animal behaviors and their underlying neural substrates. Yet, we lack principled approaches to incorporate such inductive biases into RNN models, especially in a computationally efficient manner.

We hypothesize that RNNs can develop more robust and animal-like learning of complex tasks by adopting pretraining procedures that better reflects the animal’s learning experience. Inspired by this perspective, we created a novel pretraining paradigm that explicitly trains a set of useful, basic computational skills. These subtasks are then combined via animal-matched behavioral shaping towards a target end goal (Fig. 1). The computational elements (working memory, sensory evidence integration, etc.) are themselves learned from previous experience with simpler tasks, outside of the target task context (i.e., they entail distinct loss functions separate from the target loss). We term this form of training as *kindergarten curriculum learning* (kCL), to reflect the fact that the initial learning involves basic tasks that the animal would have learned through prior training experience (i.e., kindergarten). Our approach is similar to traditional curriculum learning in that the difficulty of the task increases over time, and to meta-learning in that the curriculum tasks are assumed to share underlying computational structure with the target. It is unique in that it emphasizes a compositional view of complex tasks as a combination of simpler elements, which need to be computed in parallel and which can be pretrained with simpler means.

**Figure 1:**
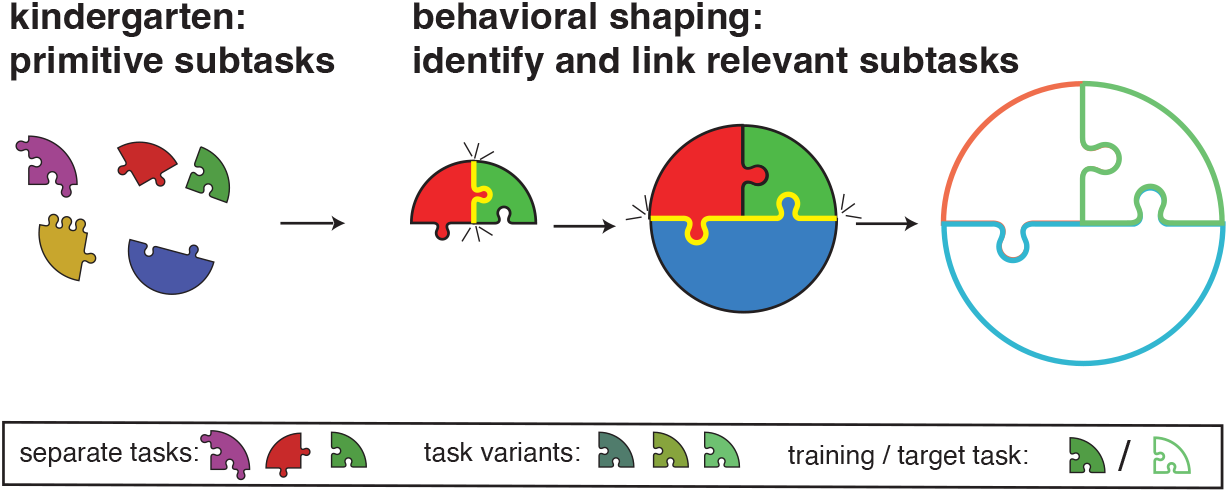
Modeling the animal’s learning experience. The target behavior or task (open circle, right) can be thought of as composed of several computational sub-elements, some of which may need to be computed in parallel. These sub-computations involve primitive skills that in the animals have likely been learned through past experience (left). Behavioral shaping guides learning by incrementally combining basic skills into more complex functions (linked puzzle pieces). For RNN training, we operationalize this idea by first training a set of fundamental sub-computations in a “kindergarten” training phase, followed by behavioral shaping which mimics animal training.

We demonstrate our general approach using a complex cognitive task previously performed in rats [19]. Formalizing the task as a Markov decision process (MDP) simplifies the problem in a way that allows us to define key qualitative features of optimal behavior in the task. We also use its solution to identify subtasks that may be relevant to include in the kindergarten curriculum. This pretraining substantially improves learning speed and target task performance compared to brute force in-task training with or without animal-like behavioral shaping. Importantly, kCL converges to computational strategies that closely match animal behavior, something not observed with a task-irrelevant kindergarten curriculum. An expansive analysis of alternative curricula demonstrates that increasing the number of relevant subtasks in the kindergarten stage increases both task performance and the strength of this match. Critically, despite a tight coupling, good performance (as measured by average reward rates in the target task) and adopting animal-like strategies (reward and context sensitivity) can be dissociated in this task, and while several pretraining approaches yielded good performance, kCL best approximated animal behavior. To determine how different pretraining strategies influence RNNs’ task solutions, we examine the dynamics of RNN activity over the course of training. We find that there is a diversity in the types of dynamical systems features present in well-trained networks (*i.e*., point attractors, saddles, and line attractors), though typically well-performing networks possess attractor dynamics for latent states with strong beliefs, and saddles for uncertain states with semi-stable beliefs. This dynamical system solution is qualitatively distinct from high performing networks not trained with kCL. Moreover, including kindergarten tasks produces networks with richer dynamics, which can be exploited for learning the target task. These observations about dynamics not only hint at the mechanisms by which previous experience impacts neural activity in living systems, but also makes explicit testable hypotheses about how neural activity is organized when rats perform the temporal wagering task.

## 3 Results

### 3.1 Behaving animals use inference in a temporal wagering task

In order to study the effects of curriculum learning on behavior and neural activity, we sought a task in which good performance would require strategies that are difficult for RNNs to learn from experience alone. Specifically, we investigated curriculum learning procedures in the context of a temporal wagering task performed by rats [19], where a simple, suboptimal strategy leads to reasonable rewards, but optimal strategies require inference of latent states over long timescales. In this task (Fig. 2a), rats initiate trials by poking into a center port, then they hear an auditory tone, the frequency of which denotes the volume of a water reward. A side port is randomly chosen as the rewarded side on each trial, indicated by an LED. On each trial, a probabilistic reward (p=0.8) is delivered with a time delay drawn from an exponential distribution *p*(*t*) = *λ*^−1^ exp(− *t/λ*) (mean delay length *λ* = 2.5*s*). The side light turns off after this delay, signaling to the rat that reward is available. If the rat chooses the other port before reward delivery, the trial is terminated (an “opt-out” trial), and the center light immediately turns on again to signal that a new trial can begin. Most trials are rewarded, but we intentionally withheld reward on a subset of trials (p=0.2), termed “catch trials,” to see how long rats were willing to wait before exercising the opt-out choice. There are 5 reward volume offers in the task (5, 10, 20, 40 and 80 *µ*L), as well as a hidden block structure that is not cued to the rats (Fig. 2b). After 40 completed trials, there is a transition from a mixed block, which contains all 5 offer values, into a high block (red) or low block (blue) that contain a subset of offers. Importantly 20 *µ*L offers are present in all blocks. Each session starts with a mixed block, randomly choosing either a high or low block, and then deterministically alternating between mixed and high/low blocks over the session. Block transitions are pseudo-probabilistic, since on some trials rats fail to maintain fixation in the center port during the auditory stimulus delivery (“violation trials”) and 40 successful trials must be completed in each block. The animal must repeat the trial following a violation. An extensive behavioral shaping protocol initially acclimates rats to the operant chamber and trial contingencies, then adjusts the probability of reward delivery before finally introducing latent block structure (see Methods).

**Figure 2:**
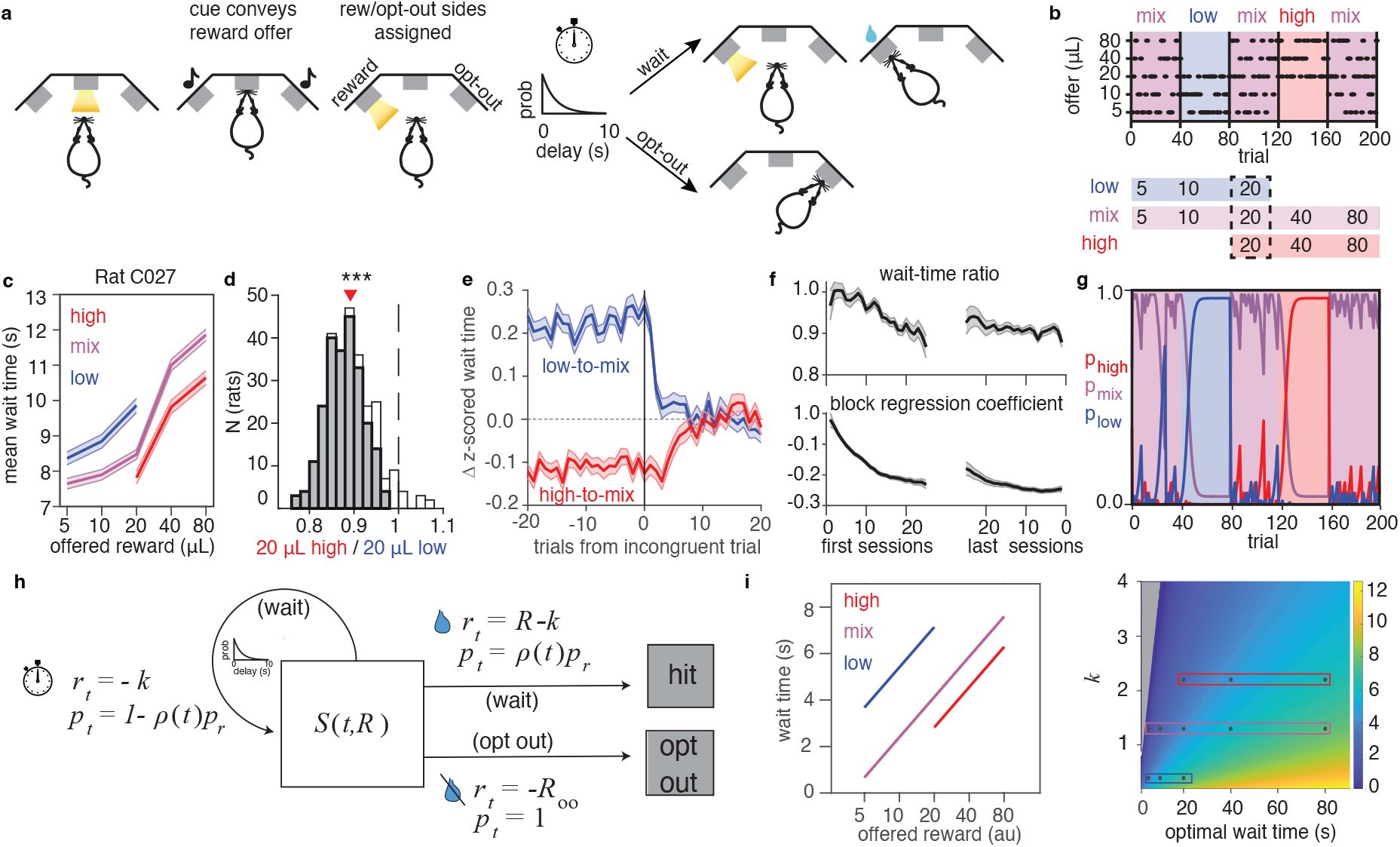
Rodent behavior in temporal wagering task. **a)** Temporal wagering task. Rats wait for known reward offers conveyed by an auditory tone, but with an uncertain reward delay. Rats may choose to wait for reward at illuminated side port until water arrives, or poke into the other port to terminate the trial. **b)** Latent block structure of task that is uncued to rats. After every 40 successful trials a transition between mixed blocks (purple) containing all potential reward offers and blocks with a subset of the highest (red) or lowest (low) offers. **c)** Example mean wait time behavior on catch trials where reward is withheld for single rat. Error bars denote s.e.m over trials. **d)** Distribution of wait time ratios of high vs. low for 20*µ*L offers on catch trials. *** : *p* = 8.26 *×* 10^−49^, (Wilcoxon signed rank test). Histogram includes rats with significantly different wait-time ratios (shaded, *p <* 0.05, rank sum test) and insignificant wait-time ratios (unshaded, *p >* 0.05, rank sum test) **e)** Trial-by-trial changes in wait times across a block transition, averaged over rats; error bars mark s.e.m., colors reflect transitions from either a high (red) or low (block) block into a mix block, aligned to first incongruent trial that violates possible offers in the current block (e.g., a 40 *µ*L offer after being in a low block). **f)** Time-course of the mean wait time ratio (top) and mean regression coefficient for block in a linear regression of wait time (bottom) for catch trials. Left: the first 20 sessions after block structure is introduced in the task; right: last 20 sessions of training. **g)** Predicted block probabilities of a Bayes-optimal model of the task. Background colors denote true block. **h)** MDP formalization of the task, showing actions (gray boxes), state-transfer probabilities *p*, and rewards *r* for each state-action pair. Rewards arrive within each trial based on the delay distribution *ρ*(*t*), incur a time penalty *k* at each timestep, as well as penalty *R*_*oo*_ for opting out. **i)** MDP-defined optimal wait times for three different wait time penalties (left). Wait times for all wait time penalties (right). Colored boxes denote *k* for optimal wait times in left plot. Gray region denotes negative wait time. In panels d-f,the mean is over rats (N=291), and error bars in e-f mark s.e.m.

On catch trials, rats are forced to exercise the opt-out option. Their wait times on these trials are sensitive to both the size of the reward offer and the latent block structure of the task (Fig. 2c). Rats wait longer for larger reward offers, and also modulate their wait times based on the reward statistics of the current block. Similar sensitivity to local reward statistics has also been observed in humans in “willingness-to-pay” paradigms [20]. Additionally, there is often an asymmetry in the magnitude of change to wait times for 20 *µL* offers when comparing low vs. high blocks: when comparing to mixed blocks, wait times in low blocks have a relatively larger shift in wait times than high blocks. To quantify behavioral sensitivity to the blocks, we calculated a ratio of wait times for 20*µ*L offers in high vs. low blocks. Wait time ratios less than 1 indicate that rats wait less time for 20 *µ*L in a high block compared to a low block. Across a large population (N=291 animals), we found that rats modulate their wait times based on the block context (Fig. 2d). Computational modeling and detailed analysis of behavior indicated that this state-dependent behavior relies on an inferential strategy, in which rats infer the most likely state based on the current offer value and their prior beliefs about the current reward block [19, 21]. This inference-based strategy accounts for aspects of the rats’ behavior, including the behavioral dynamics at state transitions (Fig. 2e). Indeed, regressing wait times against previous reward and block over training shows that the block regressor increases with the same time-course as the behavioral sensitivity to blocks, suggesting that contextual sensitivity reflects learned knowledge of block structure (Fig. 2f). We previously modeled the inferential strategy using Bayes’ rule to estimate the block probabilities based on previous rewards and knowledge of the task structure to predict the probability of being in each block on a trial-by-trial basis (Fig. 2g).

### 3.2 Optimal task behavior

Many rats exhibit linear sensitivity to log reward offers and a sensitivity to the reward blocks, so we next investigated if these behaviors reflected an optimal decision making strategy for this task. While several formalizations are possible, we chose to describe the task as a Markov Decision Process (MDP), in which agents must either wait for a reward, or opt out of the trial (Fig. 2h), similar to the approach in [22]. In this formulation, agents decide between “wait” and “opt out” actions. Waiting can either transition to the same state, or end in a terminal “hit” state where reward is delivered. Opting out deterministically ends the trial, with a reward penalty. Every time a wait action is performed, there is a time penalty *k*. The probability of being rewarded at each step is pulled from the exponential delay distribution *p*(*t*), and the probability of being rewarded on each trial is *p*_*r*_.

The MDP formulation that treats the offered reward and time within trial as the state allows for a simple closed form solution describing optimal behavior for an agent with perfect knowledge of the task statistics. Namely, for an exponential distribution *p*(*t*) = *λ*^−1^ exp(−*t/λ*), the optimal wait time *t*^*^ is log-linear in reward offer *R* as,

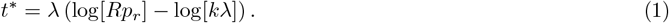

In foraging theory where agents attempt maximize their reward rates, wait time penalties are often interpreted to be related to the opportunity cost, or the average reward rate of the environment, meaning that wait time decisions should incorporate the value of potential rewards being missed out on by continuing to wait [23, 24]. In the MDP formulation here, this implies

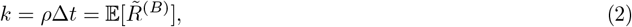

where *ρ* is the reward rate, Δ*t* is the timestep length, 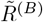 is the average reward received in a given block *B*, and the expectation is taken over the reward probability for an offer in a specific environment *B* (see also Suppl. Material). In context of the temporal wagering task, the low, mixed, and high blocks are distinct reward environments, and Eq. 1 then describes qualitative optimal behaviors in this task: rewards should be linear in log-offer, with a bias to wait longer amounts of time in reward-sparse environments (Fig. 2i). Moreover, an asymmetry in the sensitivity of wait times to high vs. low blocks, as well as a scaling of faster wait times with lower reward probabilities are also predicted (see Suppl. Material for derivations and further details).

In sum, based on the MDP formulation of our temporal wagering task, modulating wait time with linear sensitivity to the log of a reward offer is an optimal behavior. If agents further try to optimize long-term rewards in which the time penalty is the opportunity cost of the environment, then waiting longest in low blocks, followed by mixed and then high blocks is also an optimal behavior. Importantly, rats display all of these hallmarks (Figs. 2C-D), which is consistent with them using a strategy that is qualitatively similar to the MDP-defined optimal policy for this task.

### 3.3 Deep meta-reinforcement learning agents mimic rat behavior

In order to study the impact of training on behavioral performance, we modeled rats’ behavior using deep recurrent neural networks (RNNs) trained with meta-reinforcement learning (meta RL) methods that capture the ability of recurrent dynamics to support trial-by-trial learning, as opposed to relying solely on synaptic plasticity or parameter updates (Fig. 3a; LSTM architecture, all-to-all connected within each module). Meta-RL methods have recently proven useful in capturing the neural activity of frontal cortex and striatum during decision-making tasks, such as previous reward and action encoding, encoding of environmental statistics, phasic dopamine activity, and encoding of reward prediction errors [25]. Our version includes a modular architecture, with the first recurrent module implementing latent state inference (“inference” circuit), analogous to regions like orbitofrontal cortex [26–28], while the second module is responsible for action selection and representation of decision variables (“policy” layer), loosely analogous to the striatum [29].

**Figure 3:**
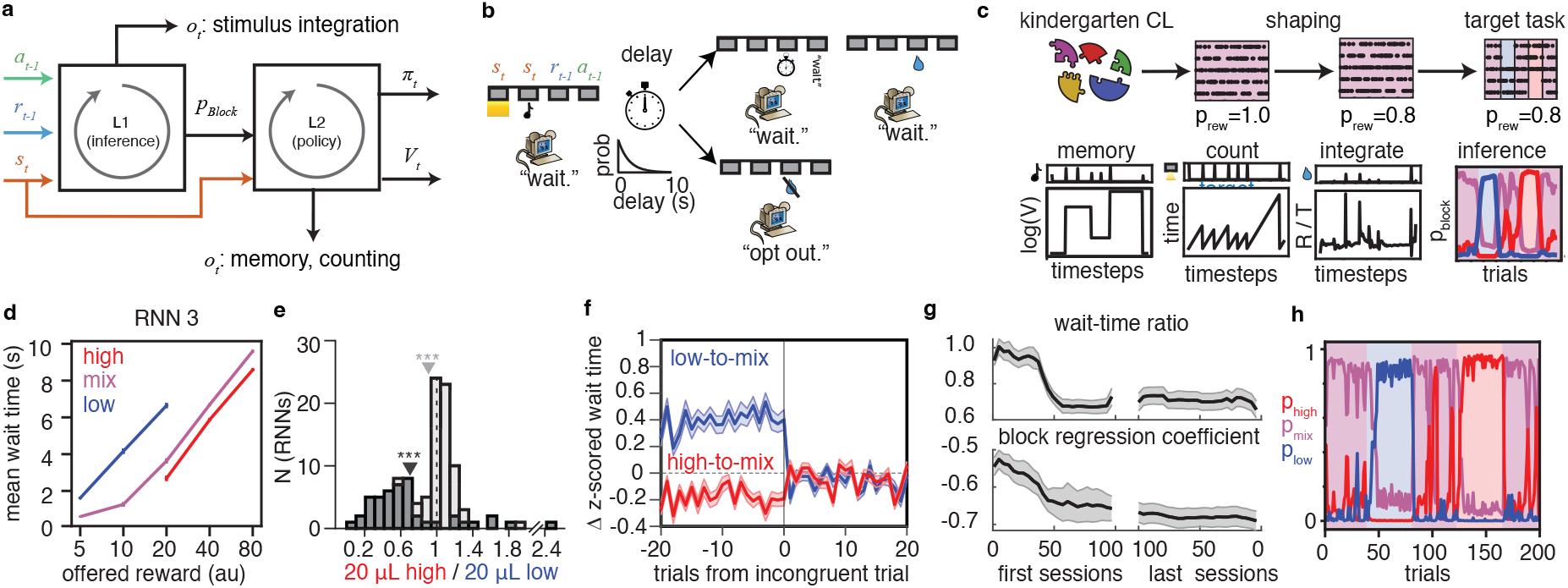
Deep meta-RL agents can learn the temporal wagering task. **a)** Schematic of deep meta-RL architecture. An LSTM-based RNN contains two modules, an inference layer that predicts latent task states, and a policy layer which outputs decisions to wait or opt out of trials. Inputs include trial stimulus information, and previous reward and action history (used in meta-RL to allow for trial-by-trial learning). The policy module outputs the decision policy, *π*_*t*_ and value function, *V*_*t*_. Additional outputs *o*_*t*_ for kindergarten tasks are used for pretraining and in-task regularization. Additionally, *p*_block_ outputs from the inference module to the policy module are trained to infer the current block. **b)** Simplified temporal wagering tasks for RNN agents. **c)** RNN training protocol, referred to as kindergarten+shaping CL. (top) kindergarten supervised learning tasks outside the context of the temporal wagering task are performed first, followed by progressively harder variants of the target wait-time task that mimics the shaping procedure performed in rats. (bottom) Normative set of kindergarten tasks suggested from MDP solution to the temporal wagering task (eq. 3): working memory, maintaining an internal estimate of time, integrating input stimuli, and inferring latent state **d).** Example mean wait time performance of RNN agent. Error bars denote s.e.m. over trials. **e)** Wait-time ratio of population of *N* = 113 RNNs. Dark bars denote *N* = 46 RNNs trained with kindergarten+shaping CL, and light bars denote other networks with simpler CL strategies (*i.e*., shaping only CL, and target task CL (see main text). Dark gray arrow is mean of kindergarten+shaping CL population, and light gray arrow is mean of entire population. ***: *p <* 0.001. kindergarten+shaping, *p* = 1 × 10^−9^; all RNNs, *p* = 3 × 10^−20^, Wilcoxon signed rank test. **f)** RNN wait-time dynamics, as in Fig 2e. **g)** Wait-time ratio (top) and regression coefficient for block in a regression of wait time (bottom) over training (see Methods). In f-g, results are the average over RNNs trained with kindergarten+shaping CL (N=46), and error bars are s.e.m over networks. **h)** *p*_block_ during the temporal wagering task for network in d. Same color convention as Fig. 2g.

We trained RNN agents on a simplified variant of the temporal wagering task (Fig. 3 b). At every time point (Δ*t* = 50ms), the agent will make a decision based upon its stochastic decision policy. The task still requires an agent to wait for known rewards with an uncertain delay drawn from an exponential delay distribution, but simplifies the task in the following ways: i) It abstracts away left- and right-sided actions to instead take a single set of actions (“wait” or “opt-out”). ii) It omits the requirement to persevere in a center port while stimulus information is provided and instead provides noise-free stimulus information to the RNN at a single time point. iii) It modifies the inter-trial interval to be variable and stochastic in length, though out of the control of the agent. These changes allow us to preserve the core decision making process in the rat task and cleanly map the wait-time task to the MDP solutions described in the previous section, while simultaneously simplifying the RNN architecture.

For pretraining, we chose the kindergarten tasks by examining the Q value for waiting during the trial, which is a key quantity that an MDP-optimal agent must calculate in order to decide when to opt out:

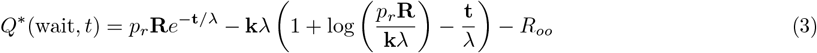

This computation involves knowledge of three key quantities (*R, t, k*), each of which point to a form of basic skill training. First, a persistent representation of *R* throughout the trial can be encouraged by working memory training. Second, knowledge of elapsed time within a trial, *t*, can be developed by training on a temporal counting task. Finally, knowledge of the opportunity cost can come via different computational strategies, either as an integrator of stimulus input that keeps track of average reward, to represent *k* through a retrospective strategy, or as a classification task that infers latent states based solely on stimulus information, which represents *k* through latent state inference. Taken together, these tasks reflect the core computations that are required for good Q value representation.

We first trained the RNN on the kindergarten tasks via individual supervised learning objectives (full description of curriculum provided in Suppl. Fig. S1), then switched to training based on the behavioral shaping protocol employed in rats: i) an easy version of the wait-time task with deterministic rewards and no block structure is performed first, followed by ii) introduction of stochastic rewards, and finally iii) introduction of the blocks in the target wait-time task. During this “shaping” phase we use an adapted version of the meta-RL loss from [25] and continue to regularize the solution with auxiliary losses corresponding to the kindergarten tasks (Fig. 3c, top). The complete schedule of training is meant to reflect the experience that rats receive before the temporal wagering task: kindergarten tasks capture preexisting cognitive capacities that the animals bring with it to the experiment (e.g., working memory, or perception of elapsed time), while the easy variants of the temporal wagering task (*i.e*., deterministic rewards and no block structure) are explicit stages of behavioral shaping for rats (see Methods).

RNNs undergoing the complete training, referred to as ‘kindergarten+shaping’ or ‘full kindergarten’ here, successfully learned to perform the temporal wagering task, exhibiting behavioral wait times on catch trials that resembled rat behavior: Agents waited longer amounts of time for larger offers, and had wait times that were sensitive to the hidden block structure (Fig. 3d). Moreover, RNNs additionally had accurate block estimates that reflected the true block (compare Fig. 3h to Fig. 2g). RNNs trained with kindergarten+shaping had strong block sensitivity across the population, as quantified by the wait time ratio, whereas RNNs without this training generally failed to become sensitive to the blocks in a manner consistent with rat behavior (Fig. 3e). Finally, when comparing the time-course of the wait time ratio and block sensitivity over training, we found that RNNs displayed similar learning dynamics to rats (Fig. 3g). Initially, RNNs do not have different wait times across blocks, nor do regressions of wait times have strong contributions from block regressors. Over training, as wait times become different across blocks for 20*µL* offers, the block regressor becomes stronger, suggesting that knowledge of block structure is driving the difference in wait times as in rats (compare to Fig. 2f). In summary, deep meta-RL agents trained with kindergarten tasks and animal-like behavioral shaping display similar qualitative behavior on the temporal wagering task as rats, have similar inferential strategies, and learn to become sensitive to latent structure that varies on long timescales in the same manner as animals.

### 3.4 Kindergarten curriculum learning supports near optimal task performance

We next tested the extent to which this structured curriculum was necessary to mimic rat behavior, and whether or not the behavior displayed by RNN agents and rats reflected an optimal decision-making strategy. We tested a variety of different CL sequences that are broadly defined using kindergarten tasks in their training, using behavioral shaping stages from the actual rat experiment, or neither type of pretraining. To keep naming conventions simple, we note that shaping is a stage of a CL sequence that reflects any experimenter-defined pretraining experienced by rats. These are RL tasks in our framework that can be identified as variations of the target task MDP, but with different hyperparameter choices (i.e., probability of reward, number of latent states). Kindergarten is a stage of a CL sequence that includes simpler tasks that can be identified from decomposing a target task. These are supervised learning tasks that are not related to the target task MDP through simple changes in hyperparameters.

We first compared the performance of networks trained with kindergarten and shaping stages to agents that experience the shaping procedure alone (Fig. 4a, orange). This second type of training reflects the final stages of shaping in the rat task. We also tested directly training on the target task without any prior training (Fig. 4a, red). In both cases, we allowed each curriculum to undergo at least the same amount of parameter updating, and often trained the networks for longer than the full set of 4 kindergarten task. We compared the reward rate during each phase of the task, and found that even when controlling for total amount of training steps, networks with training that included the kindergarten tasks outperformed both shaping training, and networks trained directly on the target task (Fig. 4b-c). To more precisely characterize these differences, we investigated key features of network behavior (Fig. 4d). In the absence of shaping, networks adopt a simple, suboptimal strategy, where agents simply wait until a timeout penalty occurs. These solutions are characterized by a low sensitivity to reward offer and context, and with opt-out rates below the true catch probability of the task. Shaping alone was able to lead to some mild sensitivity to the reward offer, though not nearly as strongly as when including kindergarten tasks. Additionally, these agents did not learn to opt out as frequently. Importantly, we found that the wait-time ratio in these networks was reversed when compared to rats and kindergarten+shaping CL, meaning that these networks waited longer for 20*µL* in high blocks. A reversed modulation of wait time by blocks reflects suboptimal behaviors in our MDP framework. Taken together, we find that RNNs and rats display the qualitative patterns of behavior that are consistent with the optimal strategy in this task, and that including kindergarten curriculum learning with shaping replicates this behavior in RNN agents.

**Figure 4:**
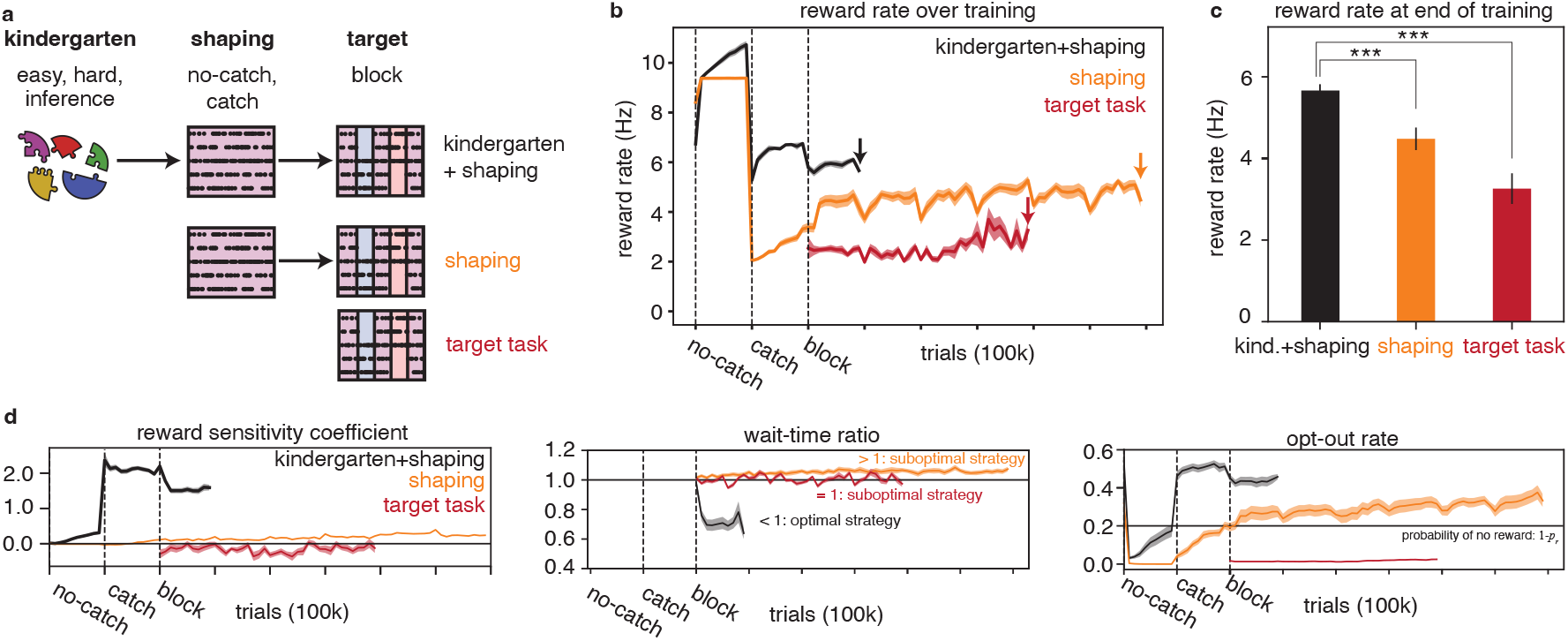
Performance of pretraining methods on the temporal wagering task. **a)** Schematic of pretraining methods, separated by kindergarten and shaping phases. The ‘kindergarten+shaping’ curriculum first trains on out-of-context supervised learning tasks (*i.e*., kindergarten tasks), followed by the final stages of behavioral shaping that occur in rat training: first training on deterministic rewards in mixed blocks only (“no-catch” training), then mixed block trials with stochastic rewards (“catch” training). Finally, blocks are introduced (“block” training). See also Suppl. Fig. S1. The “shaping” curriculum omits kindergarten tasks, and “target task” only trains on the target task with block structure. **b)** Average reward rate over training for each method. **c)** Reward at end of training stage, corresponding to colored arrows in **b**. (*** kindergarten+shaping vs. shaping: *p* = 1 *×* 10^−7^; kindergarten+shaping vs. no shaping: *p* = 5 *×* 10^−9^, rank-sum test). **d)** Evolution of different behavioral features over training. Reward sensitivity coefficient (left) is the slope of a linear fit to the wait time vs. the log of reward offer. The wait-time ratio (middle) compares average wait times for a 20 *µ*L offer in high vs. low blocks, optimally *<* 1 under the MDP formulation. Opt-out rates (right) reflect the proportion of trials that an agent chose the opt-out option, compared to the catch probability (black line). For each sub-panel in **d**, t-tests at end of training comparing kindergarten+shaping to other CL sequences were significantly different (*p <<* 0.001). **shaping** (reward sensitivity: *p* = 3 *×* 10^−25^; wait-time ratio: *p* = 3 *×* 10^−16^; opt-out rate: *p* = 9 *×* 10^−04^). **target task** (reward sensitivity: *p* = 1 *×* 10^−15^; wait-time ratio: *p* = 1 *×* 10^−5^; opt-out rate: *p* = 5 *×* 10^−27^). **b**-**d** show mean responses, with shading denoting s.e.m over networks (kindergarten+shaping: N=45, shaping: N=45, target task: N=18).

### 3.5 Full kindergarten curriculum improves on smaller or less task-matched curricula

We originally hypothesized that a normative solution derived via our MDP would identify relevant kindergarten tasks. Having shown it to be an effective curriculum, we can now ask whether pretraining on all of these tasks was actually required. Given the combinatorial explosion of possible curricula, we chose to focus on the nature and number of tasks included in the kindergarten curriculum. Additional considerations such as the ordering of tasks, and hyperparameter choices for the kindergarten tasks are briefly explored in Suppl. Fig. S2.

First, we asked if individual kindergarten tasks can achieve high-quality solutions on their own (followed by shaping). In all cases, we trained the networks for at least as much as the full set of 4 kindergarten tasks, to balance the total amount of parameter updating that could be performed. We found that, for most tasks, single task kindergarten underperformed in terms of final reward rates compared to kindergarten+shaping (aka, ‘full kindergarten’) (Fig. 5a, top), and also failed to capture its qualitative behavior (Fig. 5a, bottom). A noteworthy exception was training the memory task, which achieved equivalent reward rates. Behaviorally, the working memory only solution had comparable reward sensitivity and opt-out rates, but showed significantly weaker context sensitivity, pointing to potential differences in the computational strategy adopted to solve the task, even among high-performing solutions.

**Figure 5:**
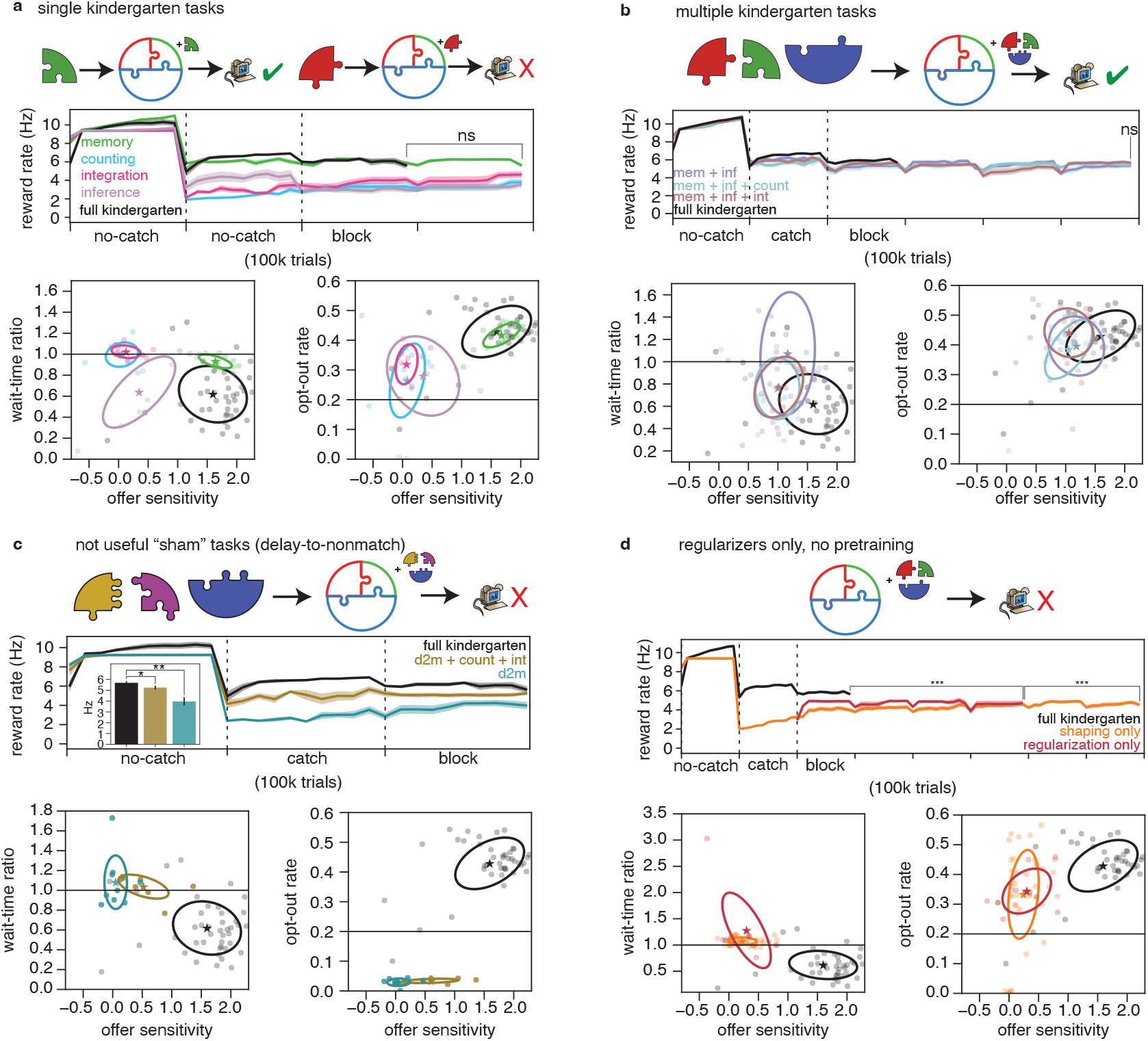
Variations of kindergarten CL. Each panel shows set of results when altering the curriculum in some form. Top: schematic of the manipulation. Large puzzle pieces denote a kindergarten task, superscript version used for regularization. Middle: reward rate over training (mean and s.e.m. over networks). Bottom: distribution of behavioral metrics for each trained RNN; stars mark population mean, and ellipses 2 std. All statistics compare full kindergarten, aka kindergarten+shaping, (N=45) vs. other CL type (rank-sum test for reward rates,t-tests for the rest. **a)** Single kindergarten tasks. **memory** (N=18, green, reward rate: *p* = 0.33; reward sensitivity: *p* = 0.53; wait-time ratio: *p* = 1 *×* 10^−4^; opt-out rate: *p* = 0.44). **counting** (N=9, blue, reward rate: *p* = 5 *×* 10^−5^; reward sensitivity: *p* = 4 *×* 10^−10^; wait-time ratio: p=3 *×* 10^−4^; opt-out rate: 3 *×* 10^−5^). **stimulus integration** (N=9, pink, reward rate: *p* = 0.002; reward sensitivity: *p* = 6 *×* 10^−10^; wait-time ratio: *p* = 2 *×* 10^−4^; opt-out rate: *p* = 0.002). **inference**: (N=9, purple, reward rate: *p* = 5 *×* 10^−5^; reward sensitivity: *p* = 2 *×* 10^−7^; wait-time ratio: *p* = 0.55; opt-out rate: *p* = 7 *×* 10^−5^). **b)** Multiple kindergarten tasks. **memory+inference** (N=18, purple, reward rate: *p* = 0.15; reward sensitivity: *p* = 0.006; wait-time ratio: *p* = 1*×* 10^−4^; opt-out rate: *p* = 0.19). **memory+inference+counting** (N=18, blue, reward rate: *p* = 0.09; reward sensitivity: *p* = 3 *×* 10^−4^; wait-time ratio: *p* = 0.08; opt-out rate: *p* = 0.17). **memory+inference+integration** (N=18, red, reward rate: *p* = 0.64; reward sensitivity: *p* = 4 10^−4^; wait-time ratio: *p* = .08; opt-out rate: *p* = 0.57). **c)** Using a computationally irrelevant delay-to-nonmatch task (d2m) **d2m** (N=9, yellow, reward rate: *p* = 2 *×* 10^−4^; reward sensitivity: *p* = 1 *×* 10^−10^; wait-time ratio: *p* = 7 *×* 10^−5^; opt-out rate: *p* = 4 *×* 10^−18^). **d2m+counting+inference** (N=9, turquoise, reward rate: *p* = 0.009; reward sensitivity: *p* = 6 *×* 10^−6^; wait-time ratio: *p* = 9 *×* 10^−5^; opt-out rate: *p* = 7 *×* 10^−18^). **d) regularization only** training performs multi-task learning with kindergarten tasks, without any CL pretraining or shaping. (N=9, red, reward rate: *p* = 0.001; reward sensitivity: *p* = 3 *×* 10^−8^; wait-time ratio: *p* = 7 *×* 10^−5^; opt-out rate: *p* = 0.02).

We reasoned that adding more kindergarten tasks might have additive effects for behavioral matching and so we studied the effect of having a 2 or 3 tasks as part of kindergarten CL (Fig. 5b). Given the already good performance of the memory-only curriculum and the good context sensitivity obtained through inference, we used these two tasks as the base pair, then optionally added a third task. All combinations retained high reward rates, but behaviorally these solutions correspond to distinct strategies, with weaker reward sensitivity and a broad range of context sensitivity (including sub-optimal reverse context sensitivity seen in shaping CL, displayed in Fig. 2d). Similarly, adding in a third task could not fully recapitulate the behavior of full kCL, though it appeared a better fit to behavior compared to the two task kindergarten curriculum. Collectively, these results demonstrate that the content of the curriculum can measurably affect behavioral strategy, even with comparable in task performance, and that the richer the (relevant) sub-computations, the closer is the trained solution to animal behavior.

We also considered whether pretraining on tasks *not* found in our normative MDP solution would lead to highquality solutions, by virtue of simply introducing task variability in training. To test this, we introduced a delay-to-nonmatch task during pretraining [30, 31], and quantified the resulting networks’ performance and decision-making strategy (Fig. 5 C). This subtask requires an agent to report if two presented stimuli that are lagged in time are the same (a match) or different (a non-match). We chose this task because it feasibly replaces the classification and working memory components learned from memory and inference kindergarten tasks. However, the essential inference component is missing from this curriculum. Hence, we hypothesized that using delay-to-nonmatch as pretraining should not lead to rat-like behavior in RNNs. We studied two versions of such pretraining using delay-to-nonmatch alone or additionally including the counting and stimulus integration tasks. Both forms of pretraining underperformed compared to kindergarten+shaping CL, and also displayed quite different behavior. This result suggests that kindergarten tasks need to contain computational sub-elements of the target task to be beneficial.

Since kindergarten tasks play multiple roles during training, we asked whether the benefits of kCL actually came from pretraining or were due to the explicit in-task regularization (Fig. 5D). Specifically, we used the full set of four kindergarten tasks but only as regularizers during in-task training (no pretraining or shaping). We found that such regularization was insufficient; it could not achieve high-quality solutions, nor could it recreate the behaviors seen in the animals, matching qualitatively the solutions obtained from behavioral shaping alone (see Suppl. Fig. S3 for average wait time curves).

We briefly studied a few additional dimensions of the curriculum structure. First, we verified that local optimum of the “simply wait” solution found through ‘target task’ CL was a persistent issue even with extended training (Suppl. Fig. S2a). Additional manipulations of the timescale of the inference task showed a temporal dependence, with short inference timescales during pretraining failing to generate strong context sensitivity (Suppl. Fig. S2b-e). This suggests that its value to the curriculum may (at least partly) stem from introducing slow modes in the RNN dynamics. Moreover, the location of the harder-to-train inference task in the sequence of subtasks also influenced the quality of the solution, consistent with the common prescription of performing simple tasks first.

Finally, one may ask how precise should the computational alignment be between kindergarten and target tasks. To answer this, we decreased the hyperparameters’ similarity between kindergarten tasks and target, while preserving the nature of the computation (different offers for memory, a larger number of latent states for inference, see Suppl. Fig. S2f). This discrepancy in the task details also meant incorporating a separate output for inference, rather than tying it to the inference-to-policy module output. We found that this generalized version of kCL can still lead to good context and wait-time sensitivity, although the resulting solutions had more nonlinear offer sensitivity, potentially due to competing computations from extra tasks (Suppl. Fig. S2f-j). This suggests that precise statistical match between kindergarten and target tasks may not be strictly needed to see benefits from kCL.

### 3.6 Kindergarten curriculum leads to distinct neural dynamics

To determine the neural substrate behind the observed learning benefits of kCL, we investigated how it changes network dynamics over training. First, we used overall magnitude of weight changes as a coarse measure of network learning (Fig. 6a). We found that large changes in network structure occurred at the beginning of each phase of training, as networks reorganize to create low-dimensional manifolds supporting structured behavior [32]. Unexpectedly, we found a substantial reorganization of the network during the kindergarten phases of training, and especially when learning to perform inference of latent states. To determine the nature of these changes in more detail, we focused on the dynamical systems structure of the solution, determined by “slow points.” The reason for this choice is the observation that storing information across time (either the reward offer within a trial or the opportunity cost across trials) is a critical aspect of the task. Slow dynamical systems features (e.g., point attractors, saddles, line attractors) provide a natural mechanism for maintenance and integration of information at long timescales [33]. Thus, identifying the computational strategy used by an RNN to solve our task translates to determining the number,type and geometry of its slow points. These can be identified numerically as local optima of the network’s kinetic energy, and their qualitative form determined by locally linearizing the dynamics around the identified slow points [34].

**Figure 6:**
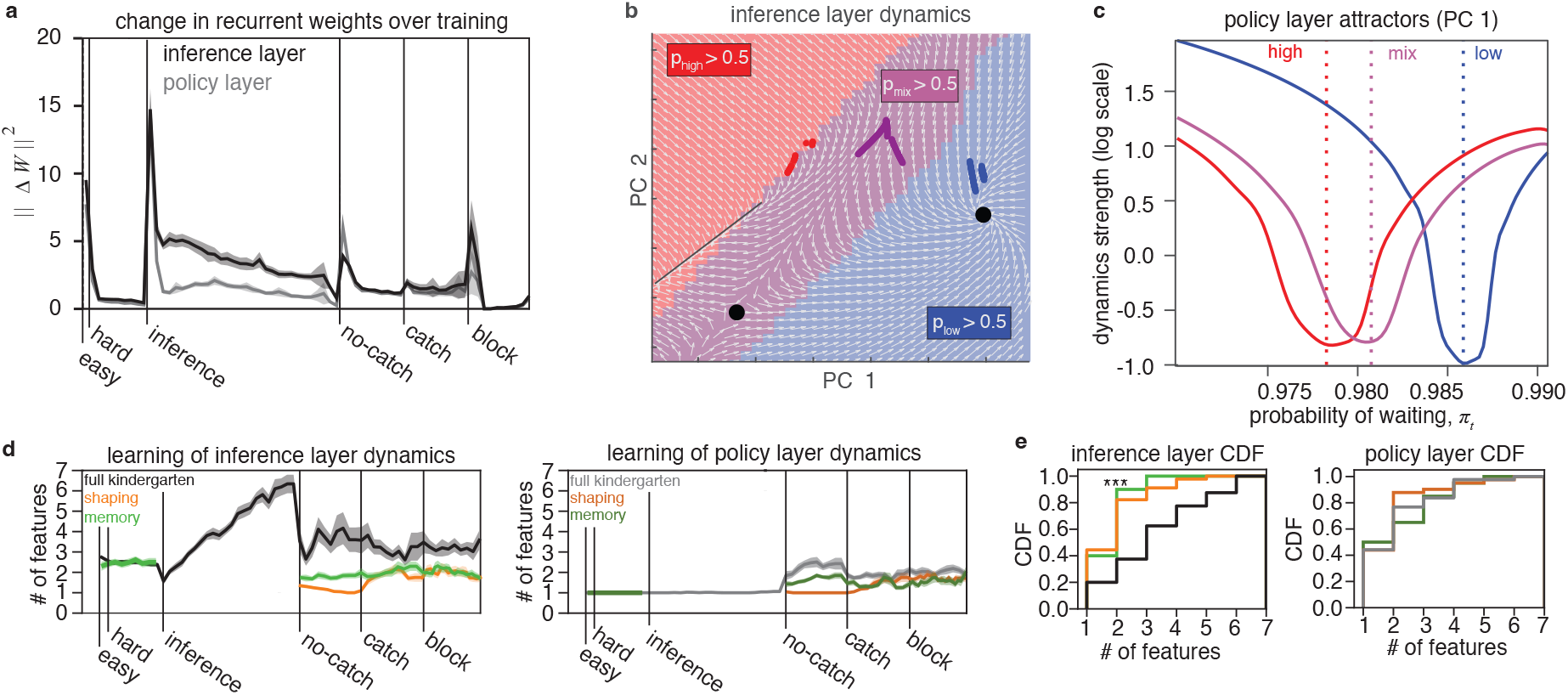
Dynamical systems features of kCL-trained networks. **a)** Change in the LSTM recurrent hidden-layer weights over training for kindergarten+shaping CL (see Methods). **b)** Dynamical gradient flow field for the inference layer of a sample network. Background color denotes region in which *p*_block_ was *>* 50% for that respective block (same color convention as in Fig. 2-3). Colored curves denote evolution of single-trial 20*µ*L offers from different blocks in which the agent ultimately opted out. Dynamical features are denoted in black (line attractor in high block, saddle in mix block, point attractor in low block). **c)** Dynamics of the policy layer along PC1, but isolated to the timepoint just before opting out for single trials from each block. PC activity was labeled with *π*_*t*_ on the x axis, and dynamics kinetic energy is plotted on log scale on y axis. **d)** Average number of dynamical systems features over training. Shaded area shows s.e.m. over networks. **e)** Cumulative distributions of the number of dynamical systems features at the end of training. A discrete KS test compared the kCL distribution at end of training to other CL distributions [35] ***: p *<* 0.001. shaping inference layer: *p* = 1 *×* 10^−4^, shaping policy layer: *p* = 0.56. memory CL inference layer: *p* = 3 *×* 10^−4^, memory CL policy layer: *p* = 0.57. Sample sizes: kindergarten+shaping N=45, shaping *N* = 45, memory N=18.

First, we investigated maintenance and updating of latent beliefs in the inference layer of kCL-trained networks (see Methods). We found that despite the common training strategy, variability in initial parameters, trial order, and the stochasticity in the policy could lead to variability in the dynamical systems solution. Nonetheless, RNNs trained by kCL generally displayed a common motif involving three slow features (Fig. 6b): two stable features (fixed point or line attractor), as well as a semi-stable feature (saddle) separating them. Among networks that learned the task in a rat-like way (29 of 45 RNNs), a large proportion (70%, or 20 of 29 RNNs) displayed this common motif. Conversely, the networks that did not learn a rat-like strategy (16 of 45 RNNs) did not display the same motif (88%, or 14 of 16 RNNs). Delineating the state space by most likely block according to the RNN’s beliefs (i.e. the inference layer’s main output), we find that the networks learn a representation that partitions state space in three contiguous sub-regions based on block confidence, with a slow feature in each (other rare motifs that still provide good wait-time behavior are documented in Suppl. Fig. S4). Given this overall structure, latent beliefs are updated by inputs from offers and rewards by pushing the network state along the direction orthogonal to the block boundaries (Suppl. Fig. S5). Importantly, the stabilizing features lie in areas of state space corresponding to low and high blocks, whereas the semi-stable feature is consistently located in the mixed-block region. Line or point attractors were equally likely in high blocks, but we invariably found a point attractor in the low block region. This overall dynamical systems structure was specific to networks trained by kCL, and almost never seen with shaping alone (2 of 47 RNNs analyzed) or other curricula (Suppl. Fig. S6a-b). Importantly, the 3-feature motif was not observed in RNNs with memory-only pretraining, (Suppl. Fig. S6c-d), despite the match in behavioral performance. This confirms the notion that different training procedures bias learned solution towards different computational strategies for performing the task. It also generates explicit testable hypotheses about the neural activity of brain regions that subserve inference (e.g., OFC)

We also analyzed the dynamics in the policy layer that support the decision to opt out. Policy layer dynamics were also variable across RNNs, but to a lesser degree than the inference layer. We often found 1-2 dynamical systems features, one of which was almost always a point attractor (41 out of 45 RNNs analyzed). We also note that the second feature was often far away from RNN activity, and only weakly influenced dynamics; in general, only one slow features was functionally relevant. We reasoned that since activity would be drawn towards the attractor, the wait-time probability at these points will be correlated with behavior. Indeed, when visualizing the location of the fixed point attractor along the first PC of policy layer activity (Fig. 6c), we found that the location of the attractors was block-specific yielding varying wait-time probabilities, and that their relative location supported asymmetric wait time behavior over blocks (Fig. 3d). We anecdotally did not observe attractor positions change across blocks in other curricula. This makes another testable prediction about neural correlates of behavior in this task, namely a block-dependent shift of attractors within the manifold of population activity which correlates with the behavioral wait-time ratios between block types.

Having seen that the structure of slow points reflects key aspects of task computation like beliefs about blocks and associated wait times, we wondered how these features may be constructed over the course of training, in particular during kindergarten (Fig. 6d). We found that kCL trained networks started building slow dynamical features from the early stages, and in particular during the inference kindergarten task. Once the temporal wagering task was introduced, these features started to get pruned from the network’s dynamic repertoire, stabilizing into the final motifs described above. This expansion and then pruning of dynamical features over training has been previously reported in other tasks [32], and might reflect a biologically relevant learning trait for tasks that rely on attractor-based dynamics. Interestingly, while the policy layer had similar numbers of dynamical features across learning procedures, the total number of dynamical features in the inference layer was curriculum-specific with kCL showing richer dynamics than sub-optimal networks trained just using shaping (Fig. 6e). Additionally, we found that kCL RNNs operated in higher dimensional spaces compared to those trained with other protocols (Suppl. Fig. S7). Overall, these results suggest that the benefits of full kindergarten curriculum training may stem from its ability to endow the network with richer dynamics that can then be exploited when finally learning the target task.

## 4 Discussion

It has been long recognized that *“Key to shaping is identifying the essential sub-components in a task.”* [2]. Here we argue that the same insight can be used to build inductive biases that steer RNNs towards learned solutions that are both computationally more efficient and a closer match to animal behavior. Our kindergarten curriculum learning procedure decomposes a target cognitive task into its prerequisite computational elements, and first trains on simple tasks that each involve one of these computational elements. Further shaping and target task training take advantage of these inductive biases to more effectively learn a good policy. We demonstrated the utility of this approach for training RNNs on a temporal wagering task, where kCL showed not only performance and training efficiency benefits but also a tighter behavioral match to the strategies adopted by rats [19, 21] compared to a range of alternative training procedures. These properties could not be replicated by traditional forms of curriculum learning employed as part of the rat shaping protocol, with computationally irrelevant kindergarten tasks, with many subsets of kindergarten tasks, or though multitask training on kindergarten tasks without pretraining. Mechanistically, the benefits of kindergarten training could be traced back to its ability of building richer dynamical systems features, particularly in the subcircuit subserving inference. Inference was also the task that most strongly reorganized network dynamics throughout training, consistent with its critical role for supporting latent context-dependent behavior.

While we found a common dynamical systems motif in many of the well-performing networks that trained with kindergarten tasks, the exact geometry of the solution was variable across networks. Moreover, different training curricula had qualitatively different dynamical systems structure despite being matched in performance. This is unlike the “universal” dynamical systems solutions sometimes found for simpler tasks, where all networks that do well solve the task in the same way and thus are dynamically isomorphic [34, 32]. Thus, this task complexity may open interesting avenues for exploring how across-animal behavioral variability can be traced back to variability in neural activity in related brain regions, in particular orbitofrontal cortex (for latent state inference) and striatum (for policy encoding). This is important in the context of our task, as there is a high degree of variability in wait-time behavior among rats [19].

Our approach traces its roots to behavioral shaping [36] but also classic curriculum learning and meta-learning. If the idea of pretraining for RNNs is not new, previous work has focused on using incremental training to expand the trainable temporal horizon of RNNs [2], or as an assay to uncover learning principles on tasks where behavior is difficult to analyze, but that could be optimally performed even without curriculum learning [37]. It is also qualitatively different from approaches that pretrain on human or animal behavior directly in a supervised manner [38, 39], which we see as complementary. In contrast, a precise design of the curriculum was critical for our task. Our approach also relates to forms of meta-learning used to account for structure learning [40], where invariant relationships in a task are stored and reused for more efficient learning. While sharing its emphasis on common computation that generalizes across tasks, kCL is distinct in its emphasis on task compositionality, and in particular in decomposing the task into non-sequential subcomputations.

The direct composition of two simple behaviors into a complex one was previously demonstrated in [41], revealed by analyzing the Q values of deep RL agents on two separate components of a navigation task. There, they found that the Q value for the target task was a sum of the constituent subtask Q values, which was also observed in mice performing the same task. In our case, the segregation of functionality into distinct subcircuits was partially enforced via the choice of architecture, which reflects known functional distinctions between orbitofrontal cortex and striatum [26–29]. Such a segregation complicates the analysis of compositionality at the level of RNN dynamics. For this, it would be useful to trace the evolution of individual dynamics features over learning, to more directly characterize how features developed at one stage of learning support later computations, in the spirit of [32].

How well would our approach generalize to other tasks? While a general answer cannot be provided using just one example, the core idea is to break the target task into computational sub-elements, some that need to be computed in parallel. This contrasts to traditional views of compositionality (inherited from hierarchical RL) where the goal is learning to stitch together a sequence of operations [42] and relies on the availability of a simplified ideal observer of the task that pointed out these key computational subelements. Admittedly, a mathematically tractable approximation of the task may not always be available or relevant, as animals can display distinctly suboptimal behavior in other tasks, such as sequential biases and lapses [43]. Nonetheless, it may be possible to intuit at least some of the needed computational elements in the kindergarten sense. In particular, kCL pretraining targeted to improve the ability to process information across long time scales, is likely to benefit other tasks with long temporal dependencies and sparse reward structure, echoing the idea that certain basic cognitive abilities might generally support the learning of complex behavior [44, 45]. Pretraining generally useful functionality into RNNs could be achieved by jointly training on many cognitive tasks [42]. Alternatively, utilizing any available (not necessarily normative) model of animals’ behavior in the target task may serve as means for identifying candidate pretraining tasks. More broadly, our work argues that modeling complex cognitive tasks in RNNs requires careful thinking about preexisting knowledge and skills that the animals bring with them to the experiment.

## Acknowledgements

We thank Laura Driscoll, Paul Glimcher, Vishwa Goudar, Owen Marschall, Kevin Miller, and Jane Wang, for helpful discussions and comments on the manuscript. We also thank Maanav Chittireddy for fixed-point characterization of shaping-trained RNNs. This work was supported by NIMH grant 1K01MH132043-01A1 (DH) and NIMH grant 1R01MH125571-01 (CMC and CS). This work was supported in part through the NYU IT High Performance Computing resources, services, and staff expertise.

## Data availability

The rat behavioral data and the statistical model of behavior were detailed in [19], published as a Zenodo database under https://doi.org/10.5281/zenodo.10031483.

## Code availability

Code used to train RNNs, analyze data, and generate figures is available at https://github.com/Savin-Lab-Code/kind_cl

## 5 Methods

### 5.1 Animal subjects and behavior

Behavioral procedures have been published in detail elsewhere [19]. Briefly, a total of 291 Long-evans rats (184 male, 107 female) between the ages of 6 and 24 months were used for this study (Rattus norvegicus). The Long-evans cohort also included ADORA2A-Cre (N =10), ChAT-Cre (N =2), DRD1-Cre (N=3), and TH-Cre (N =12). Animal use procedures were approved by the New York University Animal Welfare Committee (UAWC #2021-1120) and carried out in accordance with National Institutes of Health standards. Animals were water restricted to motivate them to perform behavioral trials. From Monday to Friday, they obtained water during behavioral training sessions, which were typically 90 minutes per day, and a subsequent ad libitum period of 20 minutes. Following training on Friday until mid-day Sunday, they received ad libitum water. Rats were weighed daily. Rats were trained in a high-throughput behavioral facility in the Constantinople lab using a computerized training protocol. Rats were trained in operant boxes with three nose poke ports. Left and right ports contained speakers to generate audio tones, and contained lick tubes to delivery water reward. All ports contained an internal LED light inside the port.

LED illumination from the center port indicated that the animal could initiate a trial by poking its nose in that port - upon trial initiation the center LED turned off. While in the center port, rats needed to maintain center fixation for a duration drawn uniformly from [0.8, 1.2] seconds. During the fixation period, a tone played from both speakers, the frequency of which indicated the volume of the offered water reward for that trial [1, 2, 4, 8, 16kHz, indicating 5, 10, 20, 40, 80 *µ*L rewards]. Following the fixation period, one of the two side LEDs was illuminated, indicating that the reward might be delivered at that port; the side was randomly chosen on each trial. This event (side LED ON) also initiated a variable and unpredictable delay period, which was randomly drawn from an exponential distribution with mean = 2.5 seconds. The reward port LED remained illuminated for the duration of the delay period, and rats were not required to maintain fixation during this period, although they tended to fixate in the reward port. When reward was available, the reward port LED turned off, and rats could collect the offered reward by nose poking in that port. The rat could also choose to terminate the trial (opt-out) at any time by nose poking in the opposite, un-illuminated side port, after which a tone was played, and new trial would immediately begin. On a proportion of trials (15-25%), the delay period would only end if the rat opted out (catch trials). If rats did not opt-out within 100s on catch trials, the trial would terminate.

The trials were self-paced: after receiving their reward or opting out, rats were free to initiate another trial immediately. However, if rats terminated center fixation prematurely, they were penalized with a white noise sound and a time out penalty (typically 2 seconds, although adjusted to individual animals). Following premature fixation breaks, the rats received the same offered reward, in order to disincentivize premature terminations for small volume offers. We introduced semi-observable, hidden-states in the task by including uncued blocks of trials with varying reward statistics: high and low blocks, which offered the highest three or lowest three rewards, respectively, and were interspersed with mixed blocks, which offered all volumes. There was a hierarchical structure to the blocks, such that high and low blocks alternated after mixed blocks (e.g., mixed-high-mixed-low, or mixed-low-mixed-high). The first block of each session was a mixed block. Blocks transitioned after 40 successfully completed trials. Because rats prematurely broke fixation on a subset of trials, in practice, block durations were variable.

#### 5.1.1 Behavioral shaping

The shaping procedure was divided into 8 stages. For stage 1, rats learned to maintain a nose poke in the center port, after which a 20*µ*L reward volume was delivered from a random illuminated side port with no delay. Initially, rats needed to maintain a 5 ms center poke. The center poke time was incremented by 1 ms following each successful trial until the center poke time reached 1s, after which the rat moved to stage 2.

Stages 2-5 progressively introduced the full set of reward volumes and corresponding auditory cues. Rats continued to receive deterministic rewards with no delay after maintaining a 1 second center poke. Each stage added one additional reward that could be selected on each trial-stage 2 added 40 *µ*L, stage 3 added 5 *µ*L, stage 4 added 80 *µ*L, and stage 5 added 10 *µ*L. Each stage progressed after 400 successfully completed trials. All subsequent stages used all 5 reward volumes.

Stage 6 introduced variable center poke times, uniformly drawn from [0.8-1.2] s. Additionally, stage 6 introduced deterministic reward delays. Initially, rewards were delivered after a 0.1s delay, which was incremented by 2 ms after each successful trial. After the rat reached delays between 0.5 and 0.8 s, the reward delay was incremented by 5 ms following successful trials. Delays between 0.8 and 1 s were incremented by 10 ms, and delays between 1 and 1.5 s were incremented by 25 ms. Rats progressed to stage 7 after reaching a reward delay of 1.5 s.

In stage 7, rats experienced variable delays, drawn from an exponential distribution with mean of 2.5 seconds. Additionally, we introduced catch trials (see above), with a catch probability of 15%. Stage 7 terminated after 250 successfully completed trials. Finally, stage 8 introduced the block structure. We additionally increased the catch probably for the first 1000 trials to 35%, to encourage the rats to learn that they could opt-out of the trial. After 1000 completed trials, the catch probability was reduced to 15-20%. All animal data in Figure 2 paper was from training stage 8. The conceptual changes that occur in Stages 7 and 8 were used in shaping of the RNNs.

### 5.2 Behavioral analyses

Behavioral sessions from rats required at least 5 catch trials to be included. Additionally, sessions were excluded if a linear regression of wait time did not have positive slope in a two-day moving-window average, or if it lacked a statistically significant positive slope coefficient for linear sensitivity on that day (F test, p *<* 0.05). Lastly, trials were excluded if the wait time was greater than two standard deviations from the mean wait time. These criteria were included primarily to exclude trials in which a rat was disengaged during the task.

For analysis of wait time ratio and regression of wait time over training, data were aggregated over groups of 2 sessions either at the beginning of block training, or the end of training (“first sessions” and “last sessions,” respectively, in Fig. 2e and Fig. 3f). The wait time ratio was calculated on these aggregated sessions, and a regression of wait time with regressors for current trial, one trial back, and current block were used to regress the wait time on that trial. For this, wait times for opt out trials were were z-scored, and an ordinal block was used (*B*_low_ = 1, *B*_mixed_ = 2, *B*_high_ = 3).

For block transition dynamics, we followed the approach in [19]. The wait times on catch trials of each rat and RNN were first z-scored separately for each volume, then the difference in these z-scored wait times were calculated for each volume, relative to the average z-scored wait time for that volume. This was calculated for each trial relative to an incongruent trial following a block transition. The change in wait time in Fig. 2e and Fig. 3f was an average of volume of these values. This approach was used to control for reward volume effects.

#### 5.2.1 Approximate Bayesian inference of block

The Bayesian observer in Fig. 2g was calculated based on methods from [19]. Briefly the posterior belief of block was calculated according to Bayes rule:

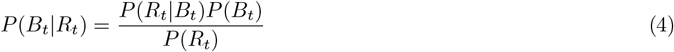

where *B*_*t*_ is the block on trial t and *R*_*t*_ is the reward on trial t. The likelihood *P* (*R*_*t*_|*B*_*t*_) is the probability of the reward for each block (1/5 for all offers in mixed blocks, and 1/3 or 0 for low and high blocks). The prior *P* (*B*_*t*_) is approximated using a posterior from the last trial as

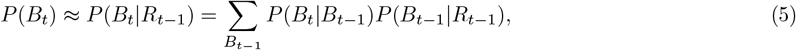

where *P* (*B*_*t*_ |*B*_*t*−1_) is referred to as the hazard rate, which incorporates knowledge of the task structure, including the block length and block transition probabilities. For example, for blocks of length *H*, the hazard rate for low blocks would be

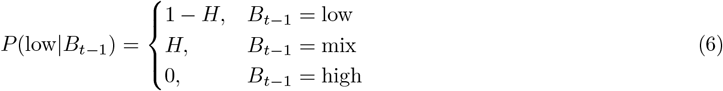

H = 1/40, to reflect the block length in this work. Extensive explanation motivating the assumption of a flat hazard rate is provided [19].

### 5.3 Details of Markov decision process (MDP) and RL formulation of task

The decision process (Fig. 2h) and RL task environment (Fig. 3b) were modeled as a simplified form of the animal task, without any left/right side choice information, requirement to persevere in the center port, or required action to initiate new trials. Formally, this was modeled as an episodic MDP with two potential actions (Wait, OptOut). The state at each point in a trial is minimally defined as a reward offer *R* and time within a trial, *t*: *S*(*R, t*). Rewards are delivered probabilistically on each trial with probability *p*_*r*_ = 0.8. Within a trial, the reward is delivered at a time that is drawn from an exponential delay distribution *p*(*t*) = *λ*^−1^ exp(−*t/λ*) (mean wait time *λ* = 2.5*s*). At every time step before reward is delivered, the agent received a small reward penalty *k* = −0.05, and when the opt-out penalty action is chosen, a reward penalty *R*_oo_ = − 2.0 is provided. The opt-out penalty was used to discourage the suboptimal strategy of instantaneously opting out. Rather than instantaneously award the reward offer or opt-out penalty, the reward was instead evenly divided out across the time bins of the inter-trial interval (ITI). ITIs were drawn from a uniform distribution from 50ms to 1s. This form of ITI better mimicked the experience in the rat task, in which animals were drinking water throughout much the ITI. It also creates longer timescale dependencies across trials, and receiving positive reward (rather than no reward) during the ITI improves value function estimates across trials. To avoid contaminating learning of the policies for Wait and OptOut, we enforced a required WaitITI action that was given unit probability of occurring during the ITI, and zero during the waiting epoch. We do not model violations in the RL formulation of the task, nor do we model the decision to begin new trials. These reflect separate computational processes that are separate from the core decision to wait or opt out.

Reward offers followed an alternating block structure, as in the rat task, with random transitions. The same alternating block structure was used, with sessions always beginning with a mixed block, and then transitioning into a low block. Note, in the rat task, there is equal probability of high or low blocks being the second block, though this difference is likely negligible to RNN behavior. Blocks were a minimum of 40 trials long, and after 40 trials block transitions occurred probabilistically, drawn from a binomial distribution with transition probability *p* = 0.5

### 5.4 Model architecture

The RL agent is comprised of a two-layer network, each with 256 LSTM units (Fig. 3 a), with relevant gates, states and inputs denoted by (*j*) for layer *j*. The computational LSTM unit of the network uses the following gates to update the hidden states *h*_*t*_ and cell states *c*_*t*_:

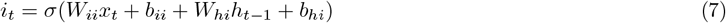

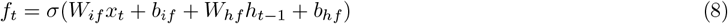

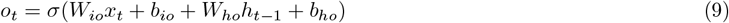

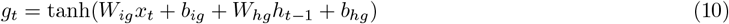

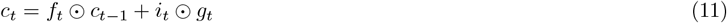

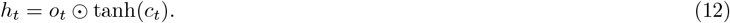

where ⊙ is the elementwise Hadamard product, *σ* is a sigmoid nonlinearity. 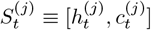 is a compact description of the activity of each layer, and successive states of the network are written here with a shorthand *S*_*t*+1_ = RNN(*S*_*t*_) or *S*_*t*+1_ = RNN(·)[(*S*_*t*_)]. Network inputs *x*_*t*_ are multidimensional, and will generally consist of external task stimuli *s*_start_ and *s*_R_, past timestep reward *r*_*t*−1_, past timestep action *a*_*t*−1_, outputs from the network *o*_*t*_, and states *S*_*t*_ from earlier LSTM layers. *s*_start_ = 1 and *s*_R_ = log(*R*) only at the start of the trial, and are otherwise zero. In contrast, the remaining inputs can in general by non-zero and time-varying over the course of an entire trial.

Specifically, the first layer (“inference” layer) receives inputs about the task stimuli of trial start and reward offer, as well as the previous reward and previous action: *x*^(inference)^ = [*s*_start,*t*_, *s*_R,*t*_, *r*_*t*−1_, *a*_*t*−1_]. The inference layer projects onto a 3-unit linear projection head that outputs the log-probability of each block 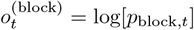, as well as a linear projection head for an auxiliary task that aims to output the average reward within a trial 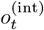. The second layer (“policy” layer) receives block probabilities from the inference layer as inputs, as well as the task stimuli for trial start and reward offer: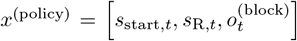. The policy layer outputs onto linear projection heads for the i) policy *π*_*t*_, ii) value of current state *V*_*t*_, iii) prediction of reward offer for a supervised memory task 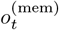, and iv) prediction of time within a task for a supervised memory task 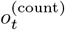 The policy *π* is a three-unit RNN with a softmax nonlinearity, corresponding to probabilities for waiting within a trial (waitTrial), opting out of a trial (OptOut), or waiting during the ITI (waitITI). During the trial, the only allowed options are waitTrial and Optout, with waitITI artificially set to low probability. During the the ITI period in which reward is delivered, the only option is waitITI. The remaining projection heads output scalar values.

Hidden and cell states of the LSTMs were initialized with random values drawn from a normal distribution (𝒩 (0, 1)). The weight parameters of the model were initialized to small values, drawn from a uniform distribution (𝒰 (− 1*/N*, 1*/N*), where *N* is the number of hidden layer units). States were reset at the end of each training epoch, where a training epoch was over all data for kindergarten costs. States were reset every 160 trials during the temporal wagering task. This reset corresponds to roughly the same timescale as progressing through all three block types, though it is not exactly aligned with block transitions.

### 5.5 Training

All RNN model training was conducted in PyTorch (v 1.8.0). For all costs, weights were updated using back-propagation through time with an Adam minimizer. Hyperparameters for training are provided in the subsequent section.

#### 5.5.1 Kindergarten training

Kindergarten tasks were comprised of supervised learning tasks: memory, counting, integration, and inference tasks. The memory, counting, and integration tasks were trained using supervised learning of a mean-squared error loss, where RNN outputs *o*_*t*_ were trained to match a target output *o*_*t*,targ_. The memory task trained the network to output the initial reward offer throughout the duration of a trial (*o*_*t*,targ_ = *s*_*R*,0_). The counting task trained networks to count time elapsed within a trial (*o*_*t*,targ_ = *t*). The integration task trained networks to calculate a running average of the ‘previous reward’ stimulus input 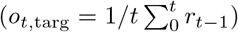. The inference task trained networks on a classification task using cross-entropy loss to categorize the latent block based on reward offer.

These tasks used similar input and trial time statistics as the target temporal wagering task. Trial durations were on the same order of magnitude as the target task, and the inputs to the RNN used the same five target values as the log-reward offer stimulus input in the RL task. The target in the integration task was based off of typical rewards received from trials in the wait-time task. Importantly, while the input statistics were similar to the target task, these kindergarten tasks are supervised learning objectives in which the actions of the RL agent have no bearing. They are fundamentally a different class of learning than the final RL-based task.

We devised a cumulative curriculum for pretraining on the four tasks by introducing tasks into training, one at a time and adding onto previous tasks (Fig. S1). We first introduced simple, single-trial variants in order of memory, counting, and integration tasks (simple kindergarten). We then increased the complexity of these tasks by extending to multiple trials in a training batch, but continuing to train on the three tasks simultaneously (hard kindergarten). Finally, we added in the inference task (inference kindergarten). We chose this ordering such that the successive tasks took more training time than previous ones.

The losses for each stage of kindergarten are the following:

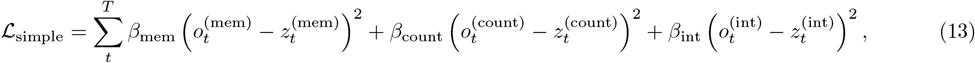

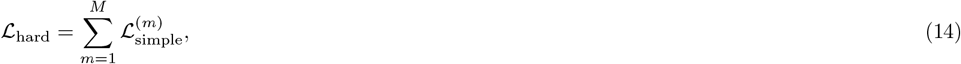

where 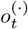 are output signals from the RNN and 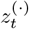 are the target outputs. 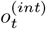 comes from the inference layer, and 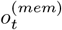 and 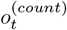 come from the policy layer. *t* denotes time within a single trial, and *m* denotes trial number. Training data for simple kindergarten was performed in a single batch, with 20 timesteps of data per sample, (*T* = 1*s*, Δ*t* = 0.05), and a batchsize of 1000 samples. Weight update occurred after every epoch over the data, and stopped when performance did not improve over 30 epochs. The second stage of kindergarten training (hard) simply expanded the time horizon of simple kindergarten, which optimized with target values from a single trial, *m*, to include multiple trials. We used *M* = 10 trials per sample, with variable trial times drawn from a uniform distribution between 1 and 5s. Training data was again a single batch, with batchsize = 1000. To maintain a single batch of data with the same amount of data per sample, each sample used 10 trials of the same durations, but in random order for each sample. Weight update was performed after each epoch over the data, and completed after a threshold of ℒ _hard_ *<* 0.001 was reached, or until 10k epochs were performed. Rather than processing each contribution to the loss cumulatively as in simple kindergarten, all 3 costs were simultaneously optimized in hard kindergarten.

Following training on simple and hard kindergarten, which were tasks with squared error losses, the final stage of kindergarten (inference) was performed. Formally, this required the outputs of the inference layer head, *p*_block_ to minimize a cross-entropy categorization loss at every time point,

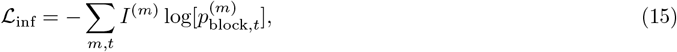

where *I*^(*m*)^ is an indicator function for the true block on each trial, *m*, taking the value *I*^(*m*)^ = 1 for the true block type, and 0 for the remaining block types. This final stage of kindergarten cumulatively optimized inference and the earlier kindergarten tasks:

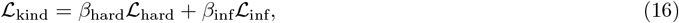

where *β*_hard_ = 1.0 and *β*_inf_ = 0.5 during kindergarten pre-training.

The sham delay-to-nonmatch task used an augmented RNN with two additional inputs to the inference layer, as well as an additional output from the policy layer. The two inputs were chosen from the the potential offers provided during the task ([5, 10, 20, 40, 80]), and there was a temporal delay between the two offers drawn from a uniform distribution 𝒰 (0.05*s*, 5*s*). There was equal probability of both inputs being the same or different on each trial. As with the other supervised learning tasks, trials were batched. The goal of the the agent was to output ‘match’ (1) if the two inputs matched, ‘non-match’ (2) if they were different, and ‘no cue’ (0) otherwise. The task was optimized using a cross-entropy loss as in eq. 15.

For the variations of the curriculum studied in Fig. 5, the ordering of tasks also proceeded as performing kindergarten tasks first, followed by shaping, and then the target task. Any task used in the pretraining stages was also used as a regularizer in during target task training, and all others were omitted as regularizers. The only exception was the ‘regularization only’ curriculum in Fig. 5d. The ordering of kindergarten tasks proceeded as performing mean-squared-error based kindergarten tasks (memory, counting, integration) before the inference task. One exception was the ‘inference first’ curriculum in Suppl. Fig. S2b-e.

#### 5.5.2 Deep meta-RL loss

Following previous work [25], the temporal wagering task described above was optimized with a deep meta-learning framework that optimized an actor-critic loss that was regularized with an entropy loss to encourage exploration, as well as the full kindergarten loss,

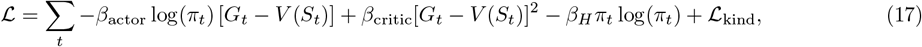

where *G*_*t*_ is the empirical, total discounted future reward

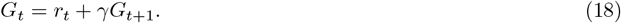

This Actor Critic approach trains networks to generate optimal decision policies *π*_*t*_ and value function estimates *V*_*t*_, but does so in a way that allows trial-by-trial learning to occur through persistent dynamical activity, as opposed to continual parameter updating of the connection weights between units. The key architectural ingredient of teaching this network to generate an internal RL procedure via its dynamics (the “meta” component) is to provide the network with explicit feedback about the past action taken, and last reward received [25]. In our two-layer architecture, this feedback is provided only to the inference layer.

### 5.6 Hyperparameters

Hyperparameters for all tasks were chosen with a combination of grid searches over groups of parameters that optimized performance and ideal behavior on the task (to balance good linear sensitivity, modest opt-out rates, low wait-time intercepts, and good inference performance), empirical searching to optimize behavior, and previous approaches as in [25]. We also utilized a heuristic approach, which was to choose regularization weights that roughly balanced the contributions of each kindergarten task in eq. (17).

#### target task

| target task |  |  |  |
| --- | --- | --- | --- |
| parameter | value | description | tuning method |
| $\alpha$ | $1e^{-4}$ | learning rate | based on [25] |
| $\gamma$ | 0.95 | discount factor | grid search |
| $\beta_{\text{actor}}$ | 1 | actor weight | - |
| $\beta_{\text{critic}}$ | 0.0005 | critic weight | grid search |
| $\beta_H$ | 0.05 | policy entropy weight | heuristic |
| $\beta_{\text{kind}}$ | 10 | hard kindergarten weight | empirical, heuristic |
| $\beta_{\text{inf}}$ | 0.5 | inference kindergarten weight | grid search |
| $\beta_{\text{d2m}}$ | 0.5 | delay-to-match weight | empirical |
| $p_r$ | 0.8 | trial reward probability | based on [19] |
| $Nt$ | 500 | timesteps per gradient step | empirical |
| $T$ | 30s | max. time before trial timeout violation | empirical |
| $\Delta t$ | 0.05 s | length of time between RNN steps | empirical |
| $T_{\text{iti}}$ | $\mathcal{N}(1, 0.15)s$ | wait time distribution between trials | empirical |
| block length | 40 trials + coin toss (p=0.5) | trials per block | based on [19] |
| state reset | 160 | trials before reset of network | empirical |

#### simple and hard kindergarten

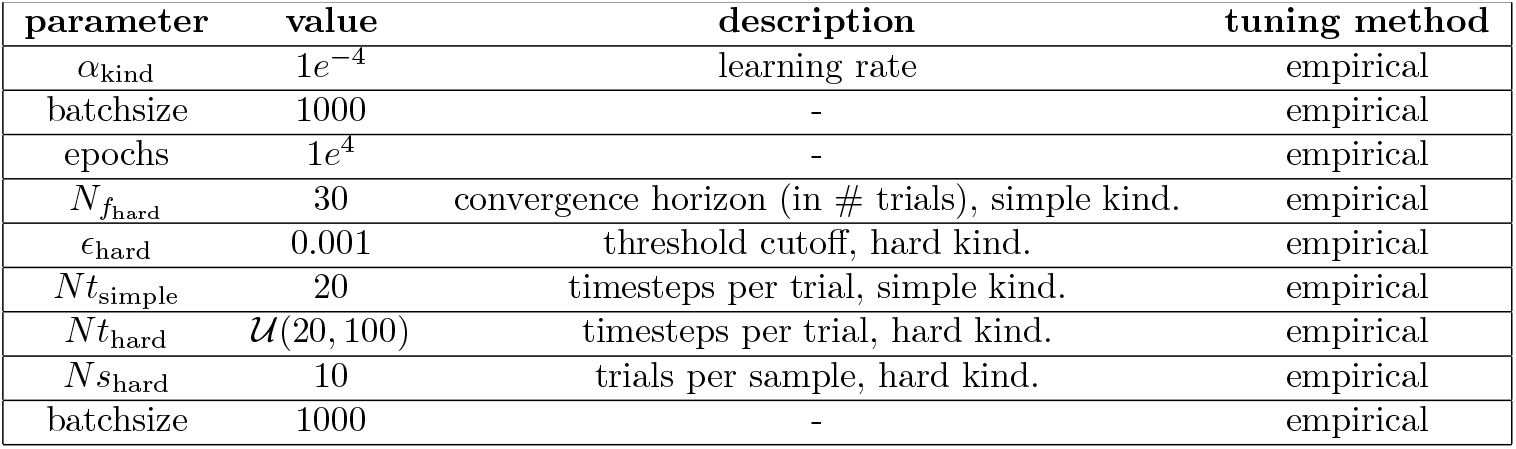

| simple and hard kindergarten |  |  |  |
| --- | --- | --- | --- |
| parameter | value | description | tuning method |
| $\alpha_{\text{kind}}$ | $1e^{-4}$ | learning rate | empirical |
| batchsize | 1000 | - | empirical |
| epochs | $1e^4$ | - | empirical |
| $N_{f_{\text{hard}}}$ | 30 | convergence horizon (in # trials), simple kind. | empirical |
| $\epsilon_{\text{hard}}$ | 0.001 | threshold cutoff, hard kind. | empirical |
| $Nt_{\text{simple}}$ | 20 | timesteps per trial, simple kind. | empirical |
| $Nt_{\text{hard}}$ | $\mathcal{U}(20, 100)$ | timesteps per trial, hard kind. | empirical |
| $Ns_{\text{hard}}$ | 10 | trials per sample, hard kind. | empirical |
| batchsize | 1000 | - | empirical |

#### inference kindergarten

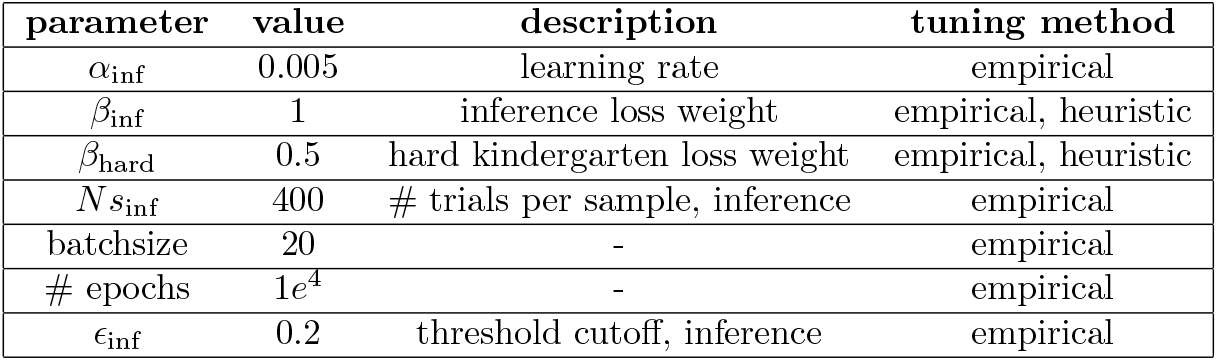

| parameter | value | description | tuning method |
| --- | --- | --- | --- |
| $\alpha_{\text{inf}}$ | 0.005 | learning rate | empirical |
| $\beta_{\text{inf}}$ | 1 | inference loss weight | empirical, heuristic |
| $\beta_{\text{hard}}$ | 0.5 | hard kindergarten loss weight | empirical, heuristic |
| $N_{s_{\text{inf}}}$ | 400 | # trials per sample, inference | empirical |
| batchsize | 20 | - | empirical |
| # epochs | $1e^4$ | - | empirical |
| $\epsilon_{\text{inf}}$ | 0.2 | threshold cutoff, inference | empirical |

#### Delay-to-match kindergarten task

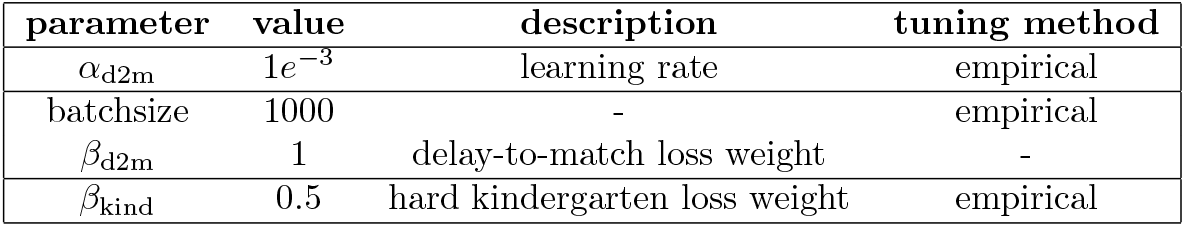

### 5.7 Network analyses

To determine which phases of training created large structural changes in the network, we calculated the change in weights Δ*W* over training. Specifically, we investigated the squared L2 norm of the difference of a concatenated recurrent weight matrix *W*≡ [*W*_*fh*_, *W*_*gh*_, *W*_*oh*_, *W*_*ih*_], over training. Different phases of training had different learning rates, so to meaningfully compare the magnitude of changes over training we sampled the network at different rates across training, proportional to their learning rates. Intuitively, this means we sampled training phases less frequently when only small parameter updates were possible (smaller learning rate), and conversely sampled more frequently in training epochs where large parameter updates were possible (larger learning rate). Specifically, the change ∥Δ*W* ∥ ^2^ in each stage is given by

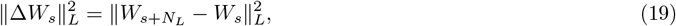

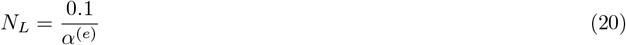

*N*_*L*_ is the number of gradient steps between samples of network weights, and was empirically chosen based on the relative learning rates *α*^(*e*)^ in each stage. For simple kindergarten, hard kindergarten, and the temporal wagering task *α*^(*e*)^ = 1*e*^−4^ (N=1000 steps). For the inference stage of kindergarten, *α*^(*e*)^ = 0.005 (N=2 steps).The wait-time task has variable numbers of gradient steps across trials, but empirically we observed that 1000 gradient steps spanned approximately 10k trials. Thus, we sampled the network every 10k trials for the temporal wagering task.

### 5.8 Reduced dynamics flow fields

A low-dimensional manifold of activity was found through conventional means by performing PCA on *S*^(*i*)^ from a session of 1000 trials from the temporal wagering task. PCA was performed separately for each layer of the network to give a reduced set of of activity 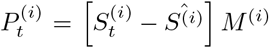, where *M* ^(*i*)^ are the principal components and 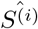 is the mean activity of network in layer *i*, respectively. Networks fully trained with kindergarten curriculum learning had ≈ 90% variance explained in the first two to three principal components. Thus, we visualized the dynamics in a two-dimensional space; however analysis of the low-dimensional dynamics was always performed in the *N* -D space that captured at least 90% variance.

Dynamical flow fields reflect the instantaneous change of the network state over time, due to network activity.

We approximate this temporal gradient with an empirical difference *F*_*t*_ across successive states as

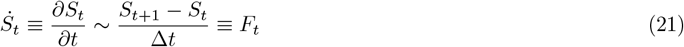

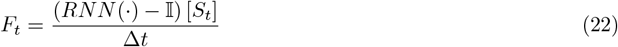

The operatore *RNN* (·) denotes propagating the state *S*_*t*_ by one timestep, and 𝕀 is the identity operator.

The flow field for behaviorally relevant low-dimensional dynamics in PC space is equivalently calculated by first projecting back into neural activity space with an inverse PCA transform, calculating *F*_*t*_, then projecting this gradient back into PC space. Assuming 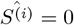 for simplicity, this dynamical gradient *F*_*P C,t*_ is given by

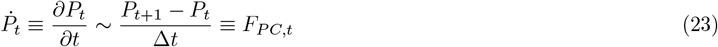

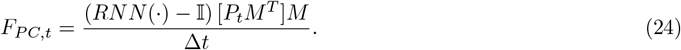

The dynamics have an intrinsic dimensionality that is typically greater than the space spanned by the variability of the network, as we noted some non-zero changes in activity in higher dimensions beyond our PCA projection (not shown). However, in practice, single-trial trajectories in these two dimensions tended to follow the flow fields of the 2-D dynamics quite well, and served as a sufficient space to analyze aspects of the dynamics that directly contribute to behavior.

Dynamics are only well-defined for static inputs. For each layer, the inputs used the wait-time epoch used in Figure 6 are *x*_(inference)_ = [*s*_start,*t*_ = 0, *s*_R,*t*_ = 0, *r*_*t*−1_ = −*k, a*_*t*−1_ = 0], and *x*_(policy)_ = [*s*_start,*t*_ = 0, *s*_R,*t*_ = 0, *o*_block_]. To calculate the reduced dynamics for the policy layer with a static input, we chose *o*_block_ values the corresponded to the block probability estimates output from the RNN during characteristic opt-out trials from each block, just before the RNN opted out.

### 5.9 Linearized dynamics

The dynamics of an RNN are in general nonlinear, and can be approximated by a Taylor series expansion around fixed point *S*_0_ as

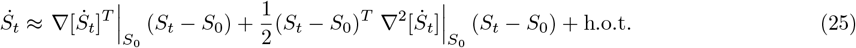

Analysis of the linearized dynamics matrix 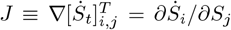 (*i.e*., Jacobian of the temporal gradient) provides a compact description of the underlying dynamical system driving behavior. The Jacobian for an LSTM only requires derivatives with respect to the cell state and hidden state. We again approximate the temporal gradient with an empirical difference, and calculate the Jacobian *J* as

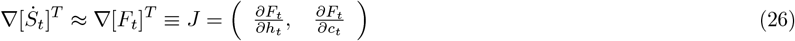

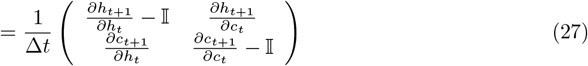

where we have used the fact that *S*_*t*+1_ = RNN(·)[(*S*_*t*_)]. The Jacobian can be written more compactly as

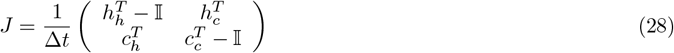

We found that the Jacobian was typically non-normal (*i.e*., when *JJ*^*T*^ ≠ *J*^*T*^ *J*), so we characterized the linearized dynamics by their spectral properties by using the Schur decomposition *J* = *WT* [46]. The Schur decomposition is analogous to a standard eigenvalue decomposition, but returns orthogonal modes, even for non-normal matrices. We characterized fixed points of the dynamics by the eigenvalues of *J* (diag(*T*)), and the Schur modes *W*. Additional details for the linearization can be found in the Supplementary Material.

### 5.10 Locating and classifying dynamical fixed points

Analysis of the dynamics for each network numerically located fixed points (*F*_*t*_ or 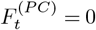); as well as “slow points” where the dynamics contained local minima, but 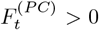. These points were located through a minimization of the kinetic energy of the system, where kinetic energy is defined by

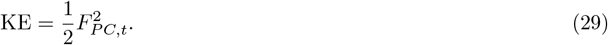

To locate fixed points in the behaviorally relevant subspace of RNN activity, we constrained the minimization of kinetic energy *F* ^(*P C*)^ in the top dimensions that explained at least 90% of variance. Thus, we searched for kinetic energy minima on the N-dimensional PCA manifold. While a fully unconstrained minimization would identify all of the fixed points of the RNN dynamics, constraining to PC space restricts our analysis to the network dynamics driving behavior. To choose initial conditions to the minimization, we used a uniform sampling for 50 points per dimension when *N* = 1 or *N* = 2. For higher dimensionality, to avoid exponential scaling issues we used a biased grid search proportional to total variance explained by each dimension. The same data and network inputs for calculating flow fields were used to calculate kinetic energy

Once minima were located, a clustering procedure (DBSCAN, scipy.clustering.DBSCAN) was performed to determine the effective number of fixed and slow points, as well as remove any outliers. The hyperparameters were min samples = 10, and epsilon. Epsilon was chosen individually for each network, as 1% of the range of support of PC1 activity. Fixed points were defined as identified minima where *KE <* 1*e*^−4^. All other minima were termed “slow points.” The network dynamics were then linearized at the identified fixed and slow points, then a Schur decomposition was performed on the Jacobian to retrieve the eigenvalues and Schur modes of the system. Dynamical systems features were categorized in the top two dimensions of PC space. Based on their eigenvalues *λ*_1_ and *λ*_2_ found by the Schur decompositions, we used the following scheme:

- *λ*_1_ *<* 0.999, *λ*_2_ *<* 0.999 : point attractor
- *λ*_1_ ∈ [0.999, 1.001] and *λ*_2_ *<* 1.001: stable line/plane attractor
- *λ*_1_ ∈ [0.999, 1.001] and *λ*_2_ *>* 1.001: unstable line
- *λ*_1_ *<* 0.999 and *λ*_2_ *>* 1.001: saddle point

In order to characterize the dynamics across RNNs, we identified a ‘common motif’ in the dynamics of the inference layer if the following features were observed: the dynamics contain a stabilizing feature (point or line attractor) in high and low-block regions of state space, as well as an unstable feature (typically a saddle) in the mixed-block region of state space. Additionally, 2-D PC space needed to contain three discrete and continuous regions of block confidence, assessed using *o*_block_. Networks were considered to have ‘rat-like’ strategies if they possessed linear sensitivity to reward offers, as well sensitivity to block context for all blocks in the same ordering as rats (longer wait times in low blocks).

## 6 Supplementary Figures

**Figure S1:**
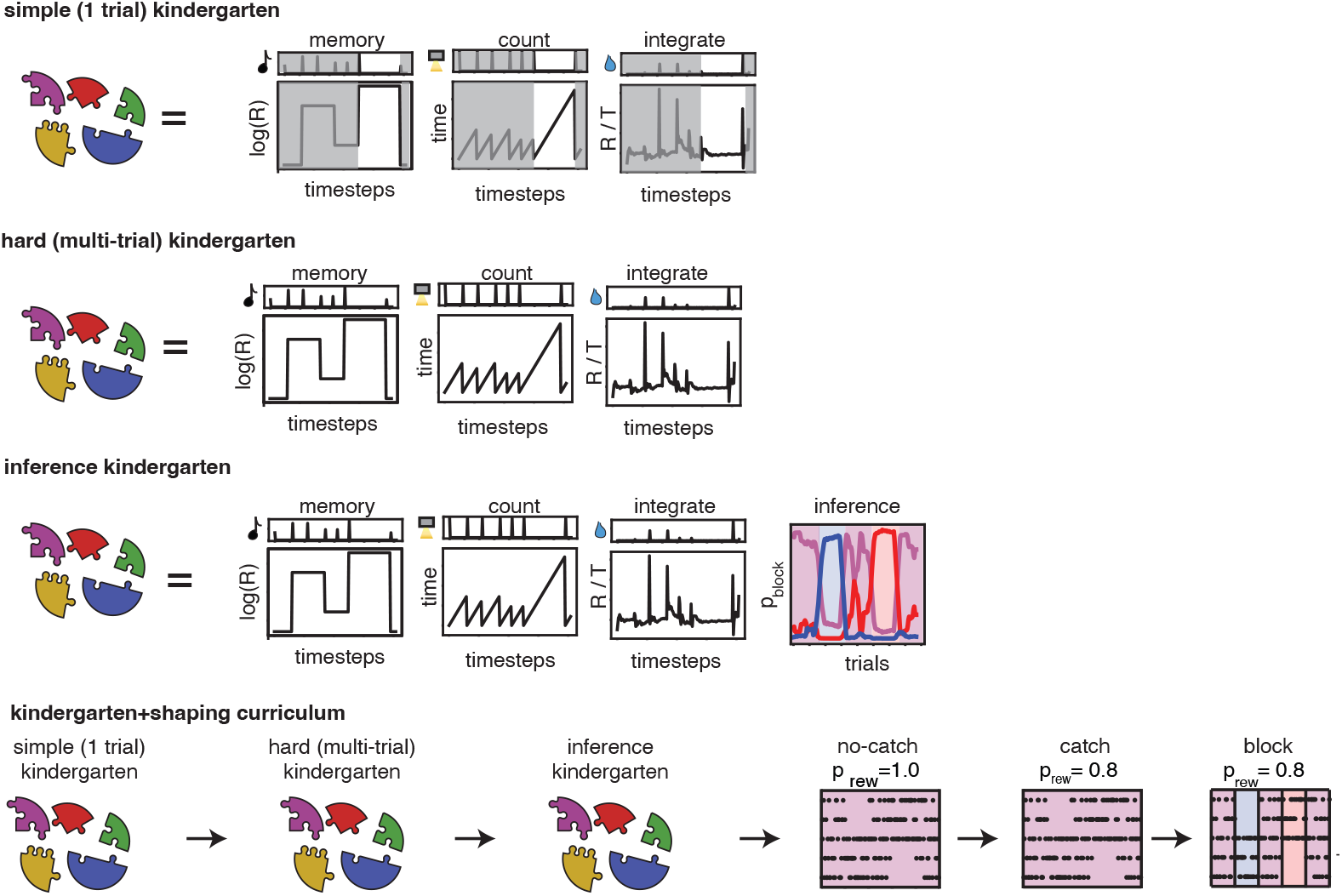
Full description of kindergarten curriculum learning. The kindergarten portion of training involves a curriculum of tasks, beginning with the memory, counting, and stimulus integration tasks over a single trial, denoted by non-masked regions (simple, 1-trial kindergarten). Following this, multiple trials are incorporated into training (Hard kindergarten). Once these tasks are performed well, the inference task is introduced (inference kindergarten). The complete kindergarten+shaping CL procedure performs these supervised tasks first, then begins training using a behavioral shaping sequence that is employed in the final stages of rat training.

**Figure S2:**
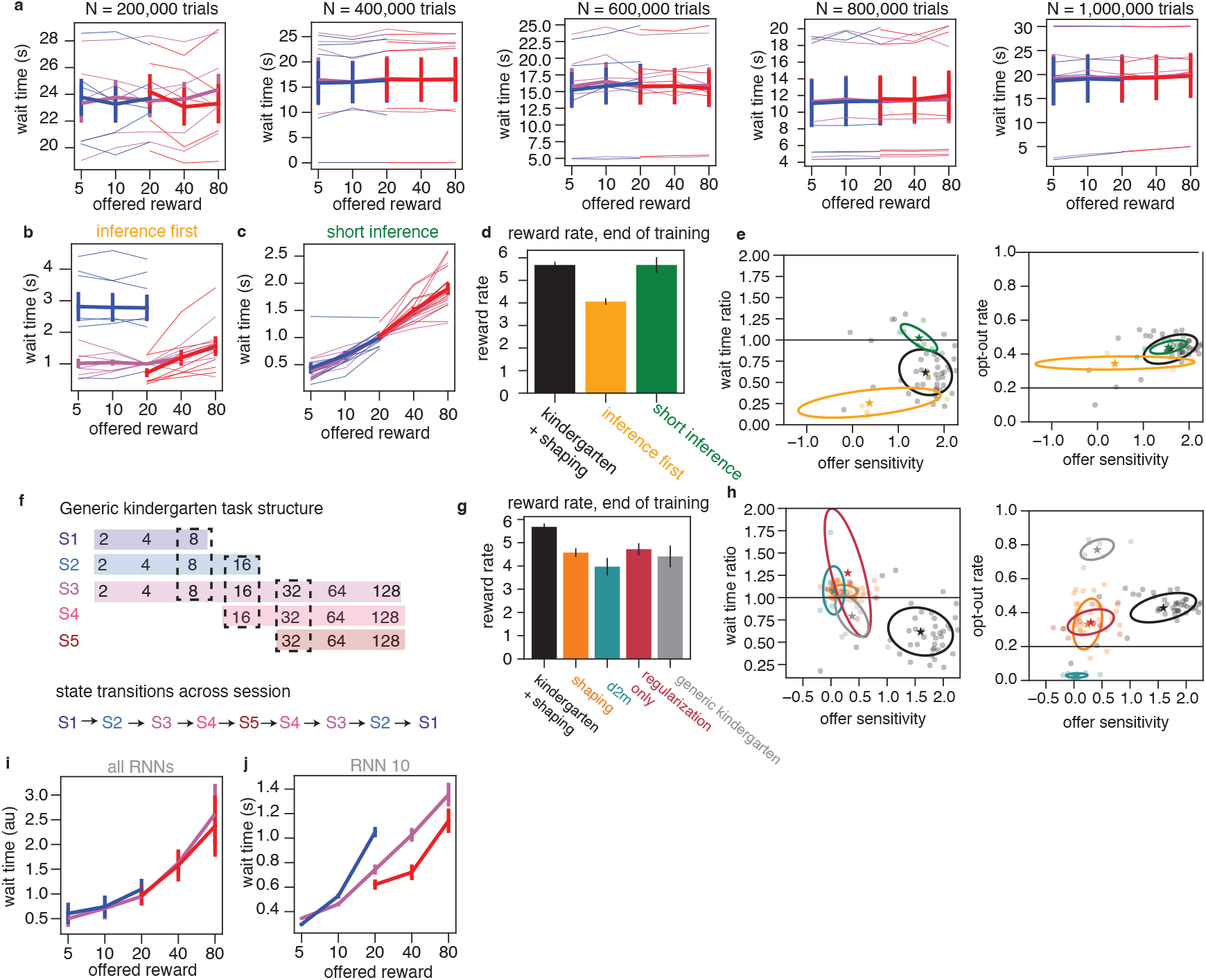
Additional CL manipulations. **a** Wait time curves for ‘target task’ CL protocol over a longer duration of training. Thin lines denote individual network wait times, and bold lines denote population mean (N=5, error bars are sem over networks). Networks appear to be stuck in suboptimal minima of “simply wait” without any reward or context sensitivity. This class of solution is not abandoned, even with training an extra order of magnitude longer on the block structure. **b-e** Manipulating the ordering and hyperparameter choices of the inference task. **b** Average and sem of wait time curves for performing inference as the first kindergarten task in the CL sequence, rather than last one. Context sensitivity is maintained, but there is a loss in performance (**d**, yellow), as well as a loss in offer sensitivity (**e**). **c** Wait time curves for changing the relative timescale of the inference task to be much shorter than the target wait-time task (N=10, error bars are sem over networks). In this form, trials had an average wait time of 0.5s, rather than 2.5 s. Networks trained on this “short inference” task maintained a high level of performance (**d**, green) as well as strong offer sensitivity **e**. However, they lose their context sensitivity, suggesting that a key feature of the inference task is to introduce slow modes into the RNN dynamics that can support long-timescale inference. **f** Schematic of a general form of inference and memory kindergarten tasks that are not directly mapped to target task statistics. Rather than doing inference over 3 states with 5 offers (*i.e*., [5,10,20,40,80]), instead these kindergarten tasks use 7 offers on a log-linear scale, across 5 states. These states transition from low-rewarding states to high-rewarding states, on offers similar but different from the target temporal wagering task. **g** Performance on this target task when training with this generic form of kindergarten CL (gray) is not as good as the statistic-matched from of kindergarten (black). These networks do retain a degree of context sensitivity and offer sensitivity, although they opt out at higher rates. **i** Average wait time curves for each RNN studied under the generic kindergarten CL, normalized to 20 *µ*L in a mixed block (error bars are sem over networks). **j** Example wait time curve from RNN trained with generic kindergarten that displays good offer and context sensitivity. Collectively, this suggests that precise statistical match between kindergarten and target tasks can improve performance, but may not be strictly needed to see benefits from kCL.

**Figure S3:**
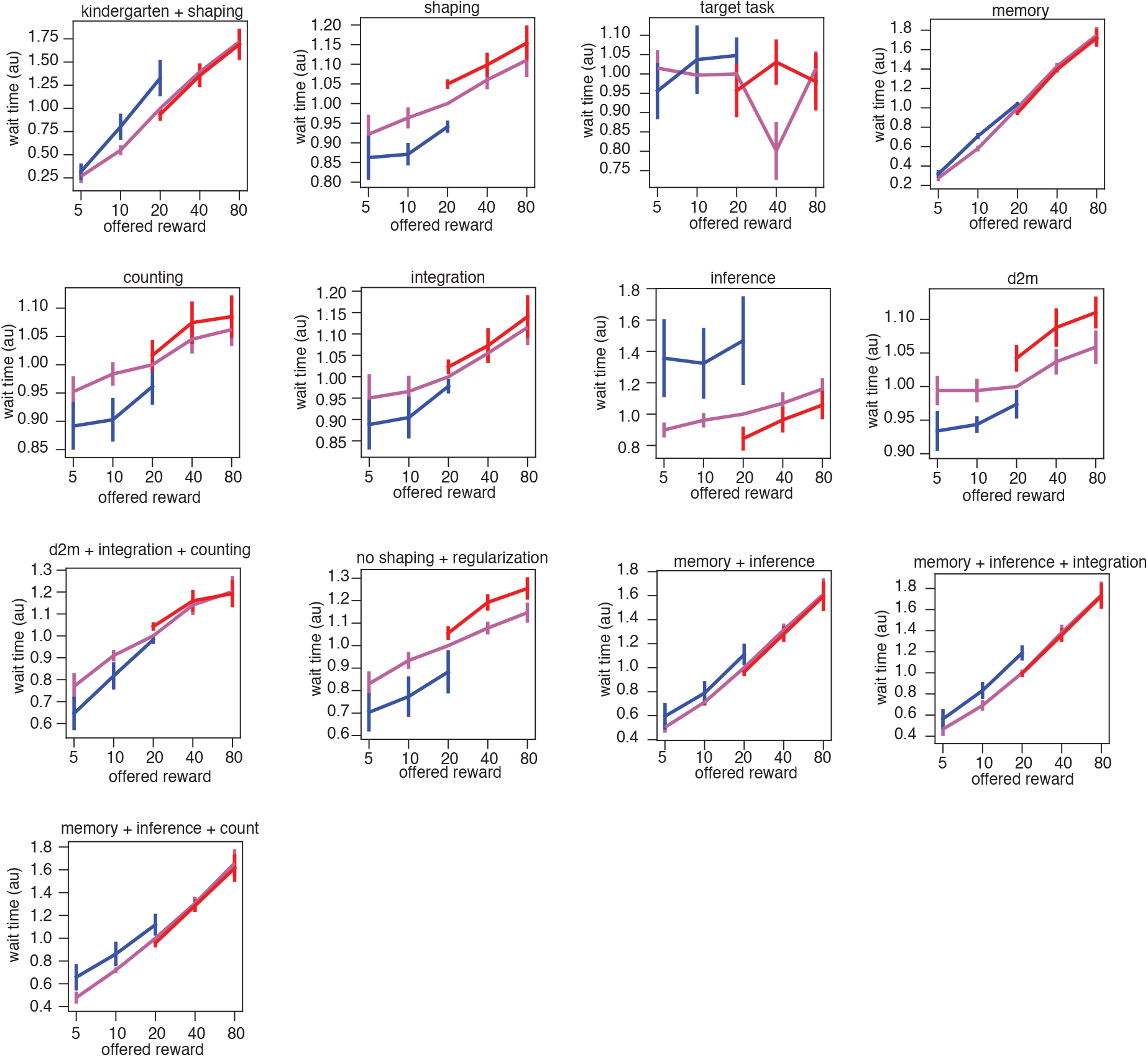
Average wait time curves across the RNN population for each CL type studied, normalized to the wait time for 20 *µ*L in a mixed block. Error bars are sem. Number of RNNs per CL type: kindergarten+shaping: N = 45; shaping: N = 45; target task: N = 20; memory: N = 20; counting: N = 10; stimulus integration: N = 10; inference: N = 10; delay-to-match (d2m): N = 10; d2m+counting+integration: N = 10; regularization only: N = 10; memory+inference: N = 20; memory+inference+integration: N = 20; memory+inference+count: N = 20.

**Figure S4:**
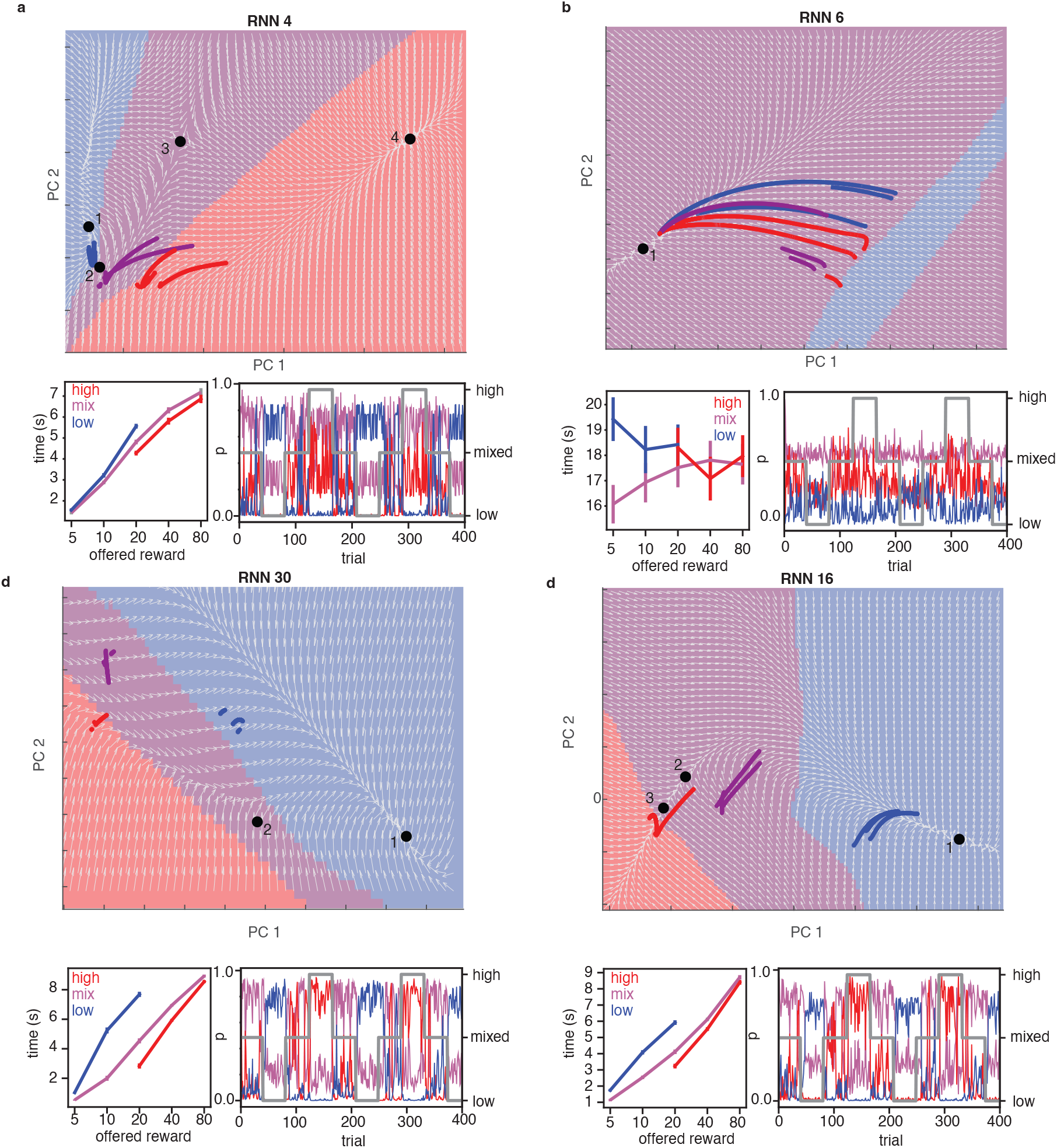
Examples of other dynamical systems motifs in top two PCS (top) found during kindergarten+shaping CL training, along with their example average wait time curves on catch trials (bottom left, error bars are s.e.m over trials) and inference performance demonstrated by plotting *p*_block_ over trials (bottom right). Located dynamical systems features are denoted by black dots and numbers. In inference performance plots, blocks are denoted by gray line. **a** sample network with same dynamical systems motif as in Fig. 6b (e.g., stable attractor dynamics in high and low blocks, saddle point in mixed block). **b** Network with poor inference performance that fails to partition state space, and has different dynamical systems features. **c-d** Alternative dynamical systems motifs supporting rat-like wait time behavior.

**Figure S5:**
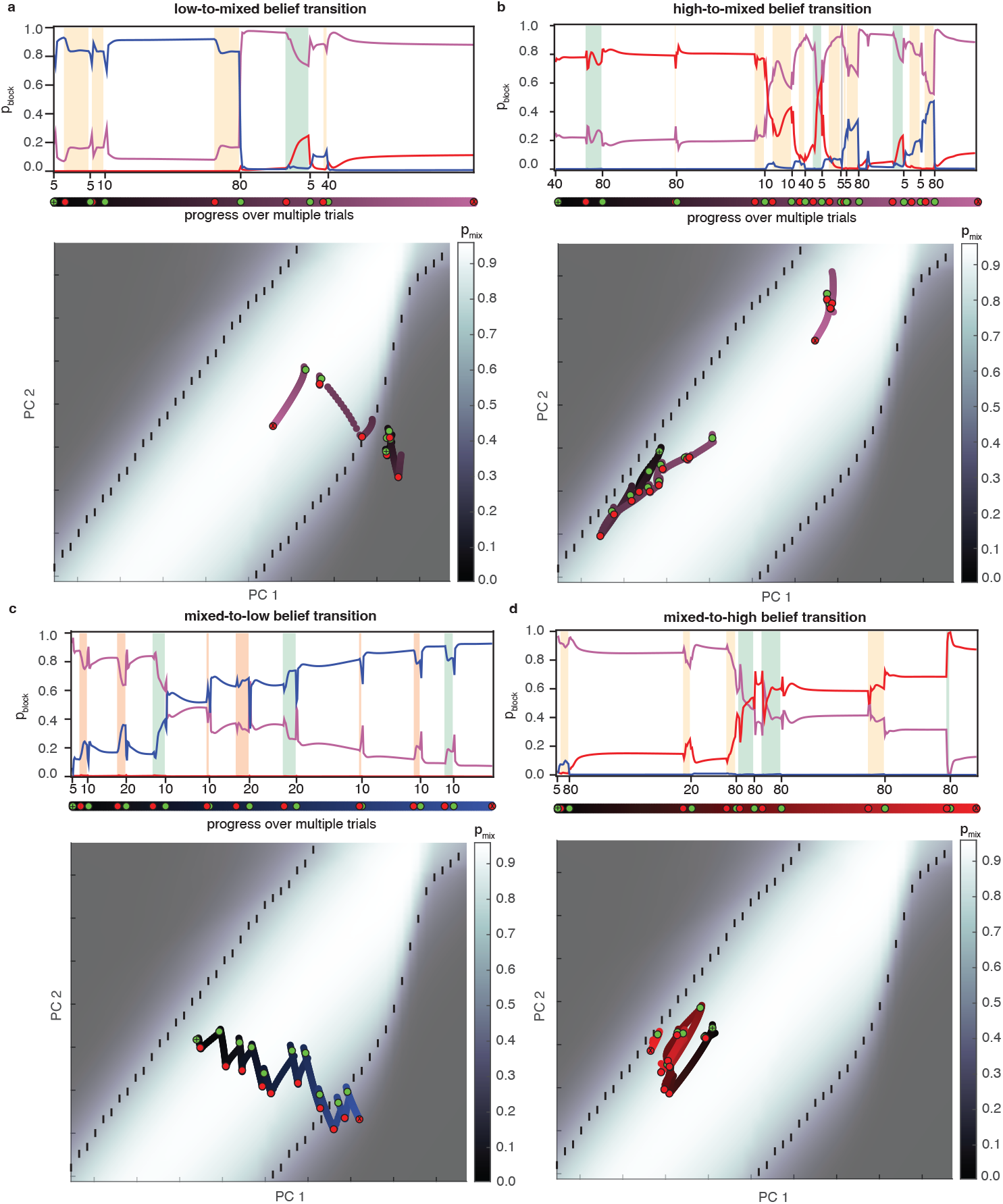
**a-d**Updating beliefs across all possible block transition types. top panels: *p*_block_ over the block transition. Trial offers are on the x axis. Colored regions denote the ITI between trials when the agent either opted out (orange) or received reward (green). White regions are the waiting epoch. Progress across multiple trials is also shown as a colored line below, with start denoted as a green dot, and the ITI as a red dot. bottom panels: analogous evolution of state activity over the trials. The background color denotes *p*_mix_, and the *p*_mix_ = 0.5 boundary is denoted with a dotted line. Trial progress uses the same convention from top plot.

**Figure S6:**
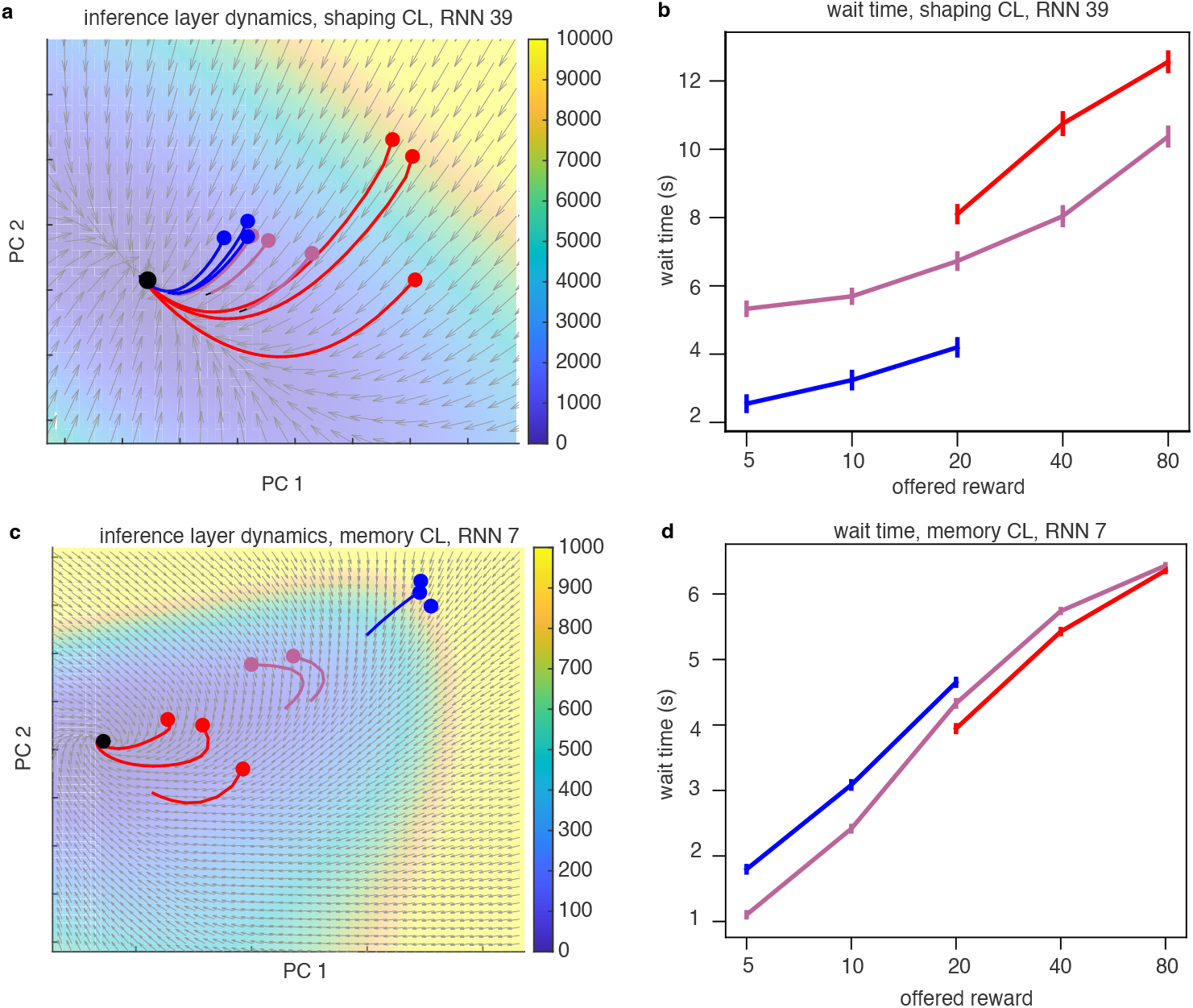
Dynamics from other CL strategies. **a** Flow field from inference layer of network trained with shaping CL, and **b** its associated wait time curve. The background colors denotes the kinetic energy. **d** Flow field from inference layer of network trained with memory CL, and **d** its associated wait time curve.

**Figure S7:**
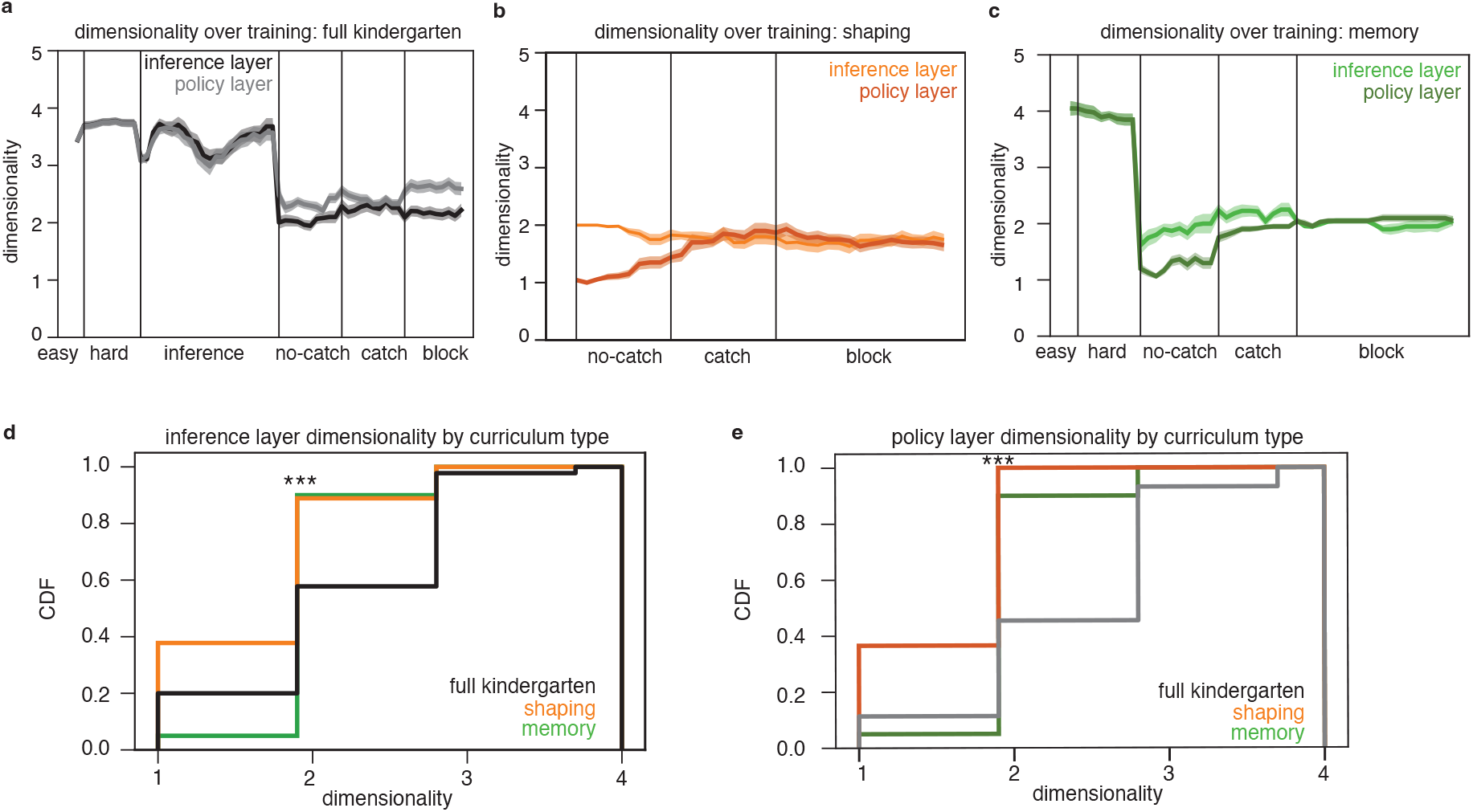
PCA dimensionality over training. **a-c** Dimensionality changes over learning for **a**) full kindergarten CL, **b**) shaping CL, and **c**) memory CL. **d-e** Distribution of PC dimensionality at end of training, for inference and policy layers. In d-e, a discrete KS test compared the kindergarten+shaping distribution at end of training to other CL distributions. ***: p *<* 0.001. shaping CL, inference layer: *p* = 1.2 *×* 10^−4^. shaping CL, policy layer: *p* = 2 *×* 10^−9^. memory CL, inference layer: *p* = 3.5 *×* 10^−4^. memory CL, policy layer: *p* = 8 *×* 10^−4^. Sample sizes: kindergarten+shaping *N* = 45, shaping *N* = 45, memory *N* = 20.

**Figure S8:**
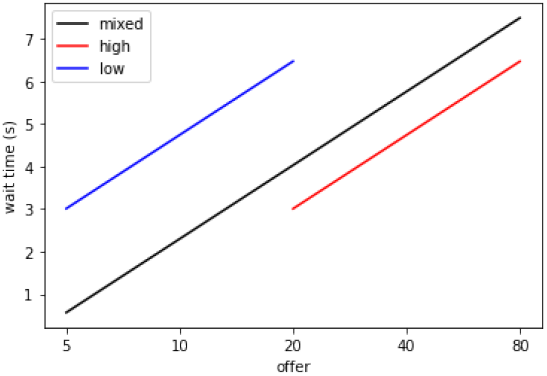
Results of SCF iteration

## 7 Supplementary Materials

### 7.1 LSTM derivatives for linearized dynamics

Here we focus on derivatives of cell and hidden state in eqs. (26-28), although full treatment of the linearization of LSTM can be found in [47]. First, an important computational identity for handling the derivative of nonlinear functions of matrix-vector products *f* (*Wx*), for *W* ∈ ℝ_*nxm*_, *x* ∈ ℝ_*mx*1_, is

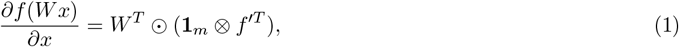

where *f* ^*′*^ = *df* (*x*)*/dx*, and **1**_*m*_ ℝ_*mx*1_. ⊗ is the Kronecker outer product. Additionally, for ∂(*x y*)*/*∂*h* we have via the chain rule,

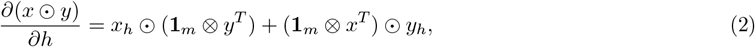

where *x*_*h*_ ≡ ∂*x/*∂*h, y*_*h*_≡ ∂*y/*∂*h*

Again, the LSTM gates are the following

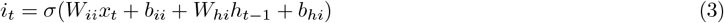

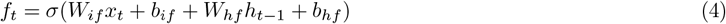

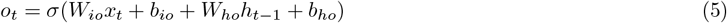

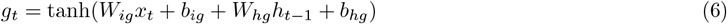

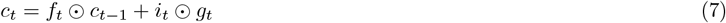

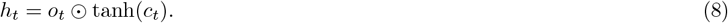

⊙ is the elementwise Hadamard product, *σ* is a sigmoid nonlinearity.

By denoting partial derivatives of gate *a*_*t*_ as ∂*a*_*t*+1_*/*∂*h*_*t*_ ≡*a*_*h*_, ∂*a*_*t*+1_*/*∂*c*_*t*_ ≡*a*_*c*_, the relevant gate derivatives for the Jacobian are the following:

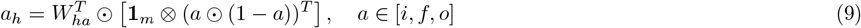

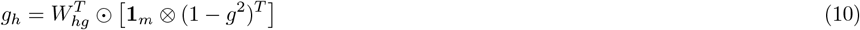

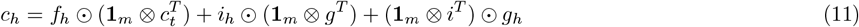

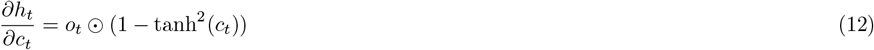

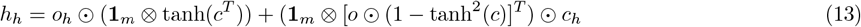

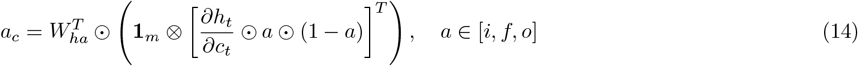

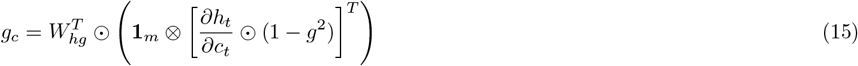

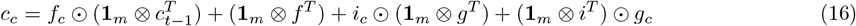

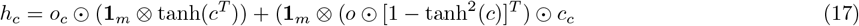

where we used eqs. (1, 2), *dσ*(*x*)*/dx* = *σ*(*x*)(1 − *σ*(*x*)), and *d* tanh(*x*)*/dx* = 1 − tanh^2^(*x*).

### 7.2 Additional MDP analysis

In this section, we provide additional details about the Markov decision process for the temporal wagering task.

#### 7.2.1 Derivation of optimal wait times

Specifically we consider the state-action values of waiting, and opting out. For an offer of size *R*, Based on the Bellman equation, these are given by [48]

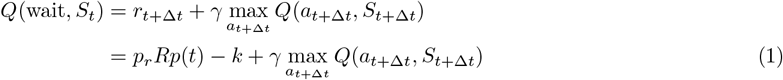

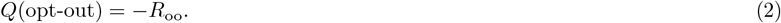

The point at which you should no longer wait is when the value of the choices are equivalent,

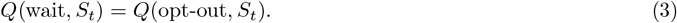

In the Bellman in eq. (1), at one step prior to this opting out timepoint, the next maximizing action should be to opt out, implying

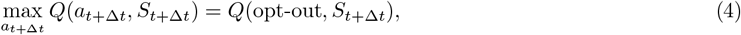

and for *γ* = 1 this yields an expression for determining optimal wait times (as in [22]):

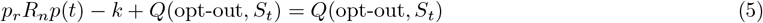

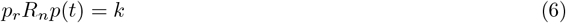

For an exponential distribution *p*(*t*) = *λ*^−1^ exp(−*t/λ*), the optimal wait time *t*^*^ is log-linear in reward offer as

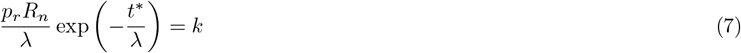

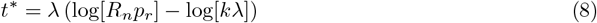

Eqs. (7-8) represent the core result that describes optimal behavior: rewards should be linear in log-offer, with an intercept that is determined by the wait time penalty. In foraging theory, wait time penalties are often interpreted to be the opportunity cost, or the average reward rate of the environment *ρ*, meaning that wait time decisions should incorporate the value of what is being missed out on by continuing to wait [23, 24]. In the MDP formulation here, we assert an approximation of the reward rate as

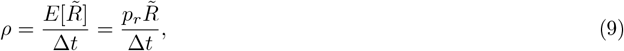

where 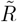 is the expected average reward from a given environment, taken separately for the low, mixed, and high blocks, respectively. If *k* is taken to be related to the opportunity cost as *k* = *ρ*Δ*t*, then shifts in the intercept of optimal wait times will be related to this log of opportunity cost:

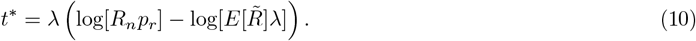

Here, 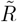 is the expected average reward from a given environment, which in the context of the temporal wagering task are taken to be low, mixed, and high blocks. The expectation is over the reward probability and delay distribution. This demonstrates that optimal behavior for a given offer *R* from an environment with higher average rewards will be to wait *less* time than if it came from an environment with lower average rewards. Intuitively, this can be thought of as being willing to wait longer for an option from a reward-sparse an environment, and less time when it happens to be the lowest option from a reward-rich environment. Simulated results from eq. (8) are shown in Figure 2 i.

#### 7.2.2 Asymmetry in wait times across blocks

As a consequence of this average reward dependence in eq. (10), there is a predicted asymmetry in the wait times across blocks in our task, where low blocks should have larger offsets from mixed blocks than compared to high blocks. Consider the difference in wait times for a given offer *R*, taken from two different environments with average rewards *R*_0_ vs. *R*_1_

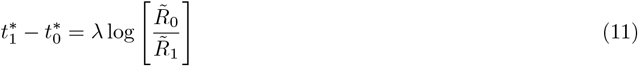

The ratio of these differences between low(L) vs. mixed(M), and high(H) vs. mixed blocks captures this asymmetry,

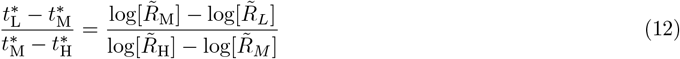

In our task, this predicts that low blocks should be approximately 2.4 times higher above the mixed block wait times, compared to the decrease for high blocks. Note, the asymmetry of RNNs and rats [19] at the level of the population is present, but substantially weaker, which is a deviation from MDP-defined optimality.

#### 7.2.3 Increased opt-out rates for higher catch probabilities

The optimal wait time in eq.(8) has a constant positive offset with the reward probability. Thus, increased catch probabilities (lower reward probabilities) are reflected in smaller wait times. Additionally, this also results in higher opt out rates: There must be at least (1− *p*_*r*_) opt outs due to catch trials; additionally, there will be a proportion of missed, rewarded trials that arrive at very long delays, given by

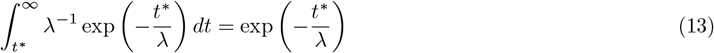

Decomposing the optimal wait time as 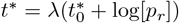 leads to the additional proportion of opted out trials as

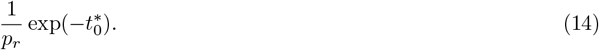

#### 7.2.4 Q values for deterministic policy

Let us consider the state-action value *Q*(wait, *R*_*n*_, *t*) ≡ *Q*(wait, *S*(*R*_*n*_, *t*)), and use a dynamic programming approach, beginning from the opt-out time of a policy, *t*_*f*_. Here we assume that *γ* = 1. This opt-out point occurs when *Q*(wait, *R*_*n*_, *t*_*f*_) = *Q*(*R*_*n*_, opt-out) = − *R*_*oo*_. Working backwards from this using the Bellman equation, you can determine one timestep back:

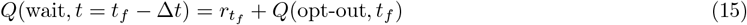

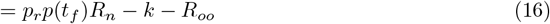

A second timestep back yields

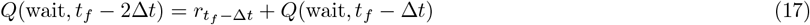

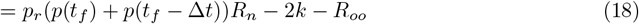

In the limit of Δ*t* → 0, this is becoming an integration back to an arbitrary time point, given as

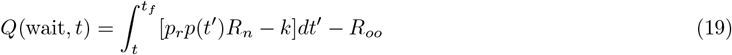

For the exponential decay distribution this yields

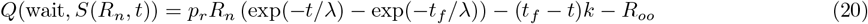

For the optimal wait time policy, then 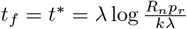, which simplifies the Q value to

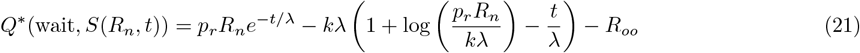

This contains an exponential decay term weighted by the reward probability, as well as a linear term and a bias.

The linear term comes from the linear penalty in the MDP.

#### 7.2.5 Reward rate for deterministic policy

Reward rates for a given policy are analytically less tractable due to unbounded exponential integrals, but numerical solutions can be found when there are minimal delivery times for reward, or for calculating differences between reward rates of different policies. The reward rate here is calculated by the expected total reward in a trial, divided by duration of a trial. Here we consider a deterministic policy that waits until *t* for reward, then opts out. On catch trials, it will simply wait until *t* and then opt out.

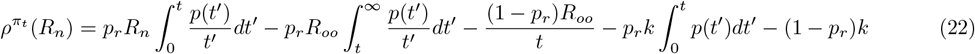

The terms above respectively accounts for i) reward arriving before time *t* and receiving reward, ii) reward arriving after *t*, and instead opting-out, iii) a catch trial, iv) the time penalty for a rewarded trial before t, and v) time penalty on a catch trial. Note, there is not a time penalty for rewards that would arrive after t, since the agent ops out, hence the presence of the integral on the wait time term for non-catch trials. The first and fourth terms are exponential integrals of the form 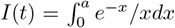, and do not have a numerical solution due to integration from *t* = 0. However, numerical solutions exist for i) integrals with a nonzero lower limit, as well as ii) differences between two integrals *I*(*t*_1_) − *I*(*t*_2_). The first case helps us understand the MVT, and the second is useful for comparing the performance different policies. We now tackle each case in turn.

If the deterministic policy has a minimal wait time *t*_0_, then the lower limit of the integrals avoids numerical instability issues, and the reward rate can be calculated numerically.

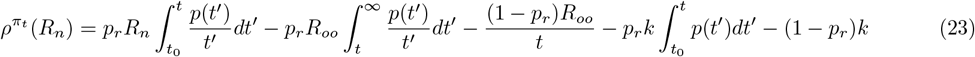

Interpreting *t*_0_ is difficult, but a loose interpretation in the rat task could be that this is the minimal time for reward to be delivered. We can also get numerically tractable integrals by comparing the reward rates between two policies that have wait different times *t*_2_ *> t*_1_:

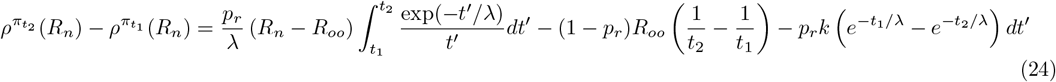

#### 7.2.6 POMDP formulation: unknown time penalty

Linear sensitivity is optimal in the case of one, unknown state with associated opportunity cost, but in the full rat task, there are multiple blocks, each with separate opportunity costs. When *k* is an unknown quantity, a POMPDP formulation would augment the state of the system to incorporate the belief about each block occurring, *p*_block_: *S*(*p*_*B*_, *n, R*_*n*_, *t*). Optimal wait times would try to optimize for expected opportunity cost, *k*_*i*_

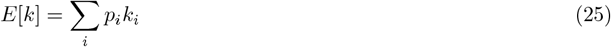

Substituting this into eq. 1 gives the optimal wait time in the presence of unknown blocks:

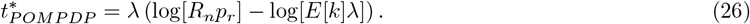

The interpretation of this result is that an optimal agent would not infer being in a single block and then choose a wait time based upon that time penalty, which would be a heuristic to optimal decision making. Nor would it average wait times over the possible wait times for each block. Instead, it would wait based upon the probability-weighted average of the possible time penalties across blocks. In the case that beliefs about blocks are very high, the heuristic of inferring a single block would have strong correspondence, but instances in which beliefs are weak about any particular block (middle of mixed blocks, or block transitions to low or high) would be able to dissociate a heuristic strategy from an optimal one.

#### 7.2.7 Optimal MVT-driven decision making: inducing an effective MDP with self-consistent reward rate and optimal policy equations

The time penalty *k* was defined in the RNN implementation to be a smaller negative reward; however, optimizing for an actor loss that aims to maximize the agent’s reward rate induces a new, effective MDP where solutions follow the Marginal Value Theorem (MVT), and *k* is then interpreted as the opportunity cost. How would this arise? Algorithmically this would occur by replacing *k* with *ρ*Δ*t* as the following:

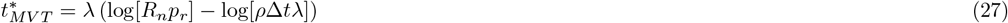

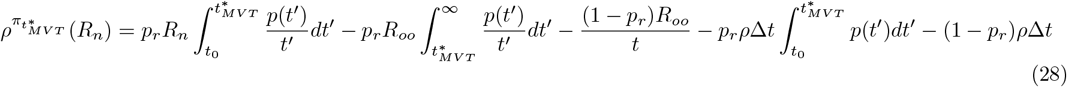

Eqs. (27-28) are known as self-consistent equations for 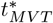 and 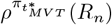, meaning that they are a set of nonlinear equations that must be they must simultaneously satisfied. Self consistent equations can be solved by iteration, where an initial guess at *ρ* can be used to calculate *t*, which can be used to solve for a new reward rate, and so forth.

Results of an iterative update of eqs. (27-28) are shown in Figures 7.2.7. One important note is that the wait times can be negative unless the lower limit *t*_0_ of the exponential integrals is set to be very low, in this case *t*_0_ = 0.01. The motivation here is not to show the absolute optimal wait times, but rather the trends among blocks. It also suggests an iterative approach for developing a strategy that incorporates the marginal value theorem: Take some guess at the reward rate of the environment and then base your wait times from that. Based on the rewards from that policy, calculate a new reward rate and repeat the process of following a new set of optimal wait times.

